# The Thermodynamics of Biomolecular CO_2_ Capture: Disentangling Equilibria in Amino-Acid-based Systems

**DOI:** 10.64898/2026.07.23.740333

**Authors:** Nicolai C. Petersen, Yang Yang, Jan S. Nowak, Jiwoong Lee, Peter Westh, Daniel E. Otzen

**Author notes:** To whom correspondence should be addressed at (D.E.O) or (P.W.).

## Abstract

Amino acids and peptides are promising building blocks for aqueous biomolecular CO_2_ capture systems, yet the coupled thermodynamics governing carbamate formation, proton transfer, carbonate speciation, and hydration remain difficult to resolve experimentally. Here, we establish isothermal titration calorimetry (ITC) as a quantitative platform for characterizing these coupled processes by integrating calorimetry with pH titrations, NMR spectroscopy, and a mechanistic thermodynamic model. Using L-lysine, L-arginine, and a series of Lys- and Arg-containing peptides, global fitting of ITC thermograms yielded thermodynamic parameters describing protonation and carbamate formation that accurately reproduced independent pH titrations and NMR-derived speciation. The analysis revealed that the characteristic biphasic calorimetric response originates from the coupled carbonate–amine equilibrium network and buffer collapse rather than carbamate saturation. Lys formed α-, ε-, and α,ε-dicarbamates and exhibited more favorable apparent carbamate thermodynamics than Arg with the ε-carbamate lying among the most favorable carbamate-forming amine sites reported for aqueous amines. Model-guided exploration of the fitted thermodynamic landscape further demonstrated that maximizing total CO_2_ retention, amine-mediated capture, and carbamate formation are distinct optimization problems governed by different combinations of pH, temperature, and CO_2_ loading. Extension to systematically spaced Lys-containing peptides showed that inter-amine separation alone does not control carbamate stability, highlighting the dominant role of the local thermodynamic environment in biomolecular CO_2_ capture. This work establishes ITC as a powerful experimental approach for extracting CO_2_–amine thermodynamics and provides a predictive framework for the rational design and optimization of amino acid-, peptide-, and protein-based carbon capture systems.

## Introduction

Current strategies to capture CO_2_ from air (global levels now at 0.04%) or flue gases (CO_2_ content typically 3-15%) are hampered by a “Catch 22” challenge: high affinity is needed to capture and concentrate CO_2_ from its natural low levels but high affinity leads to a very stable CO_2_–sorbent complex, making it energetically costly to release CO_2_ for storage or utilization^1–3^. It is essential to develop systems which tackle this challenge. This is also a major problem in amine-based scrubbing, where the extreme reaction conditions also lead to equipment corrosion, degradation of capture agents, toxicity and solvent loss through evaporation^4^. One option is to make tunable systems which switch between high- and low-affinity states based on simple and easily adjustable external parameters such as pH shifts or a moderate change in temperature^5^. Such systems include proteins and peptides, whose biological origin make them potentially available in unlimited amounts based on cheap and sustainable growth media (*e.g.* from waste streams)^6,7^. For this reason alone, they constitute attractive options as CO_2_ absorbents *if* we can develop protein-based capture strategies.

Yet there is surprisingly little experimental insight into what drives CO_2_ binding to peptides and proteins. Structural studies have focused on enzymes coupling CO_2_ to nucleophiles such as water (carbonic anhydrase)^8^, or organic molecules such as enolates undergoing carboxylation^9^, but from the perspective of interaction, these cases are obscured by the close link between binding and catalysis. Covalent carbamate formation between CO_2_ and amines is often invoked as the dominant mode of interaction in such systems^10^. This occurs continually *in vivo* in the reaction between CO_2_ and the N-terminus of hemoglobin in the blood as part of the transport of oxygen^11^. The process may be reversed when environmental changes shift the equilibrium away from carbamate formation, for example through changes in solvent conditions such as pH or through alterations in the local environment around the amino group that reduce carbamate stabilization^12^. In aqueous environments, however, CO_2_ participates in a network of coupled equilibria involving hydration, carbonate formation, and proton transfer^13,14^. These processes generate bicarbonate and carbonate species whose distributions are governed by pH and solution composition, complicating the interpretation of CO_2_ reactivity in terms of discrete amine-related molecular interactions.

To provide mechanistic insight into amino acids as CO_2_ sorbents and begin establishing principles for biomolecular capture design, it is first necessary to understand how CO_2_ interacts with the amino acid constituents of peptides and proteins. Nucleophilic amines on Lys and Arg residues can react with CO_2_ to form carbamate species, however this pathway competes directly with carbonate formation and protonation equilibria^7,15,16^. Under some conditions, carbamates are unstable and the captured CO_2_ is better described as a part of zwitterionic species of the type R-NH_2_^+^COO^-^, or fully dissociated bicarbonate accompanied by a protonated amine (*i.e.* R-NH_3_^+^and HCO_3_^-^)^17^. Consequently, CO_2_ interactions with amine groups must be described as a network of coupled equilibria, in which carbamate formation represents only one of several competing pathways **(Supplementary Figure S1)**^18^. This highlights an important principle for interpreting and optimizing amino-acid-based CO_2_ sorbents while additionally establishing a conceptual basis for future design of peptide motifs with tunable CO_2_ interactions. Given that the local environment of a Lys or Arg residue influences their pKa values and therefore the availability of deprotonated amines^19^, CO_2_ capture and speciation are expected to depend strongly on local molecular context through its effects on protonation state, carbamate stability, and partitioning between carbamate and inorganic carbon species^20^.

Here we focus on the interactions between CO_2_ and L-Lys and L-Arg residues in a simple chemical context: as free amino acids (**Figure S2)** and small dynamic peptides (**Table S1)**. This approach enables direct examination of the intrinsic chemistry governing CO_2_–amino acid interactions, decoupled from higher-order structural effects. Isothermal titration calorimetry (ITC) was employed as the primary analytical technique because of its unique ability to resolve linked equilibria through direct detection of the enthalpic response to small perturbations in sample composition during CO_2_ addition. In contrast to methods that primarily report total concentrations (e.g. spectroscopy), ITC captures the combined thermodynamic contributions from hydration, carbonate formation, proton transfer, and carbamate formation, making it particularly suited for studying coupled CO_2_–amine chemistry. Conversely, ITC provides little to no information on structure and reaction pathways. To address this, the reaction was additionally monitored by NMR spectroscopy and pH titrations. Finally, to combine these complementary datasets, we developed a titration simulator coupled to a thermodynamic equilibrium model that quantitatively describes the interconnected CO_2_–amine reaction network and resolves individual species contributions during titration.

By integrating calorimetric, spectroscopic, and potentiometric data, this work establishes a unified framework for interpreting CO_2_–amino acid interactions in complex aqueous environments (**Figure 1**). We show that the observed calorimetric behavior is dominated by proton-coupled carbonate chemistry, while carbamate formation, although directly observed, does not control the overall thermodynamic response. Apparent binding-like behavior instead emerges from the coupling of multiple equilibria within the CO_2_–amine system and suggests that capture behavior is governed primarily by protonation, speciation, and local molecular environment.

**Figure 1.**
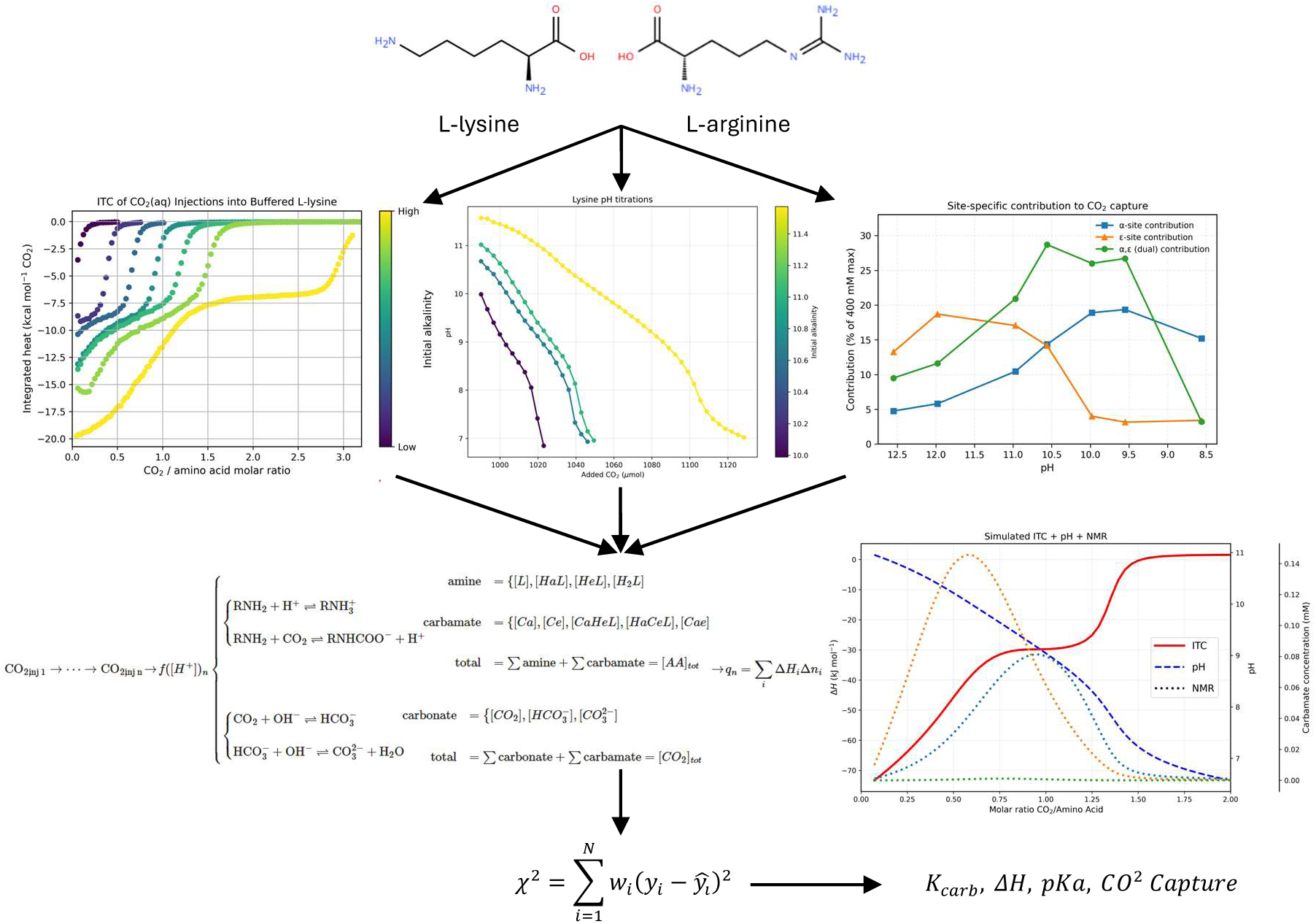
- Integrated experimental and thermodynamic framework for resolving CO_2_ speciation in amino-acid-based capture systems. CO_₂_ uptake by amino-acid-based systems was investigated by combining ITC, pH titration, NMR-derived speciation, and equilibrium modeling. Chemical structures of Lys and Arg illustrate the relevant amine-containing sites capable of participating in protonation and carbamate formation. ITC titrations of CO_₂_(aq) into buffered amino-acd show biphasic heat responses. Parallel pH titrations reveal the associated acidification during CO_₂_ addition and provide an independent constraint on the proton-coupled equilibria. NMR-derived site-specific carbamate populations show the changing contributions of α-carbamate, ε-carbamate, and α,ε-dicarbamate species to carbamate-bound CO_₂_ as a function of pH. These experimental observables were interpreted using a thermodynamic equilibrium model. The model simulates heat release, pH evolution, and carbamate populations during progressive CO_2_ loading, enabling separation of apparent binding-like calorimetric transitions from the underlying coupled speciation processes. Through a least squares fitting algorithm, thermodynamics of CO_₂_ capture can be determined and overall capture capacity can be obtained.

## Results

### Biphasic Calorimetry Reflects Intrinsic CO_2_ Reactivity

To evaluate the suitability of isothermal titration calorimetry (ITC) for probing CO_2_ interactions with amino acids, a CO_2_-saturated aqueous solution was titrated into unbuffered L-Lys and L-Arg and compared to triethylamine (TEA), a tertiary amine lacking the ability for carbamate formation, and water control (**Figure 2a**). As no external pH adjustment was applied, the experiments were performed at the intrinsic pH of the amino acid solutions in Milli-Q water, which was approximately pH 9.4–9.6 under the experimental conditions. A clear biphasic exothermic signal can be seen in both the Lys and Arg experiments while TEA and water produced only a small initial endothermic heat. While a clear signal, reminiscent of a two-site binding mechanism^21^, exists for both amino acids, it is unlikely the two available amines on Lys (α and ε-NH_3_^+^) and the α-amine and the guanidinium on Arg are responsible, as significant amine deprotonation is not expected in each system at their intrinsic pH (Lys: pH ∼9.4, Arg: pH ∼9.6). This prompted the question of what reactions of the CO_2_-amine reaction network were responsible for the enthalpic signal.

**Figure 2.**
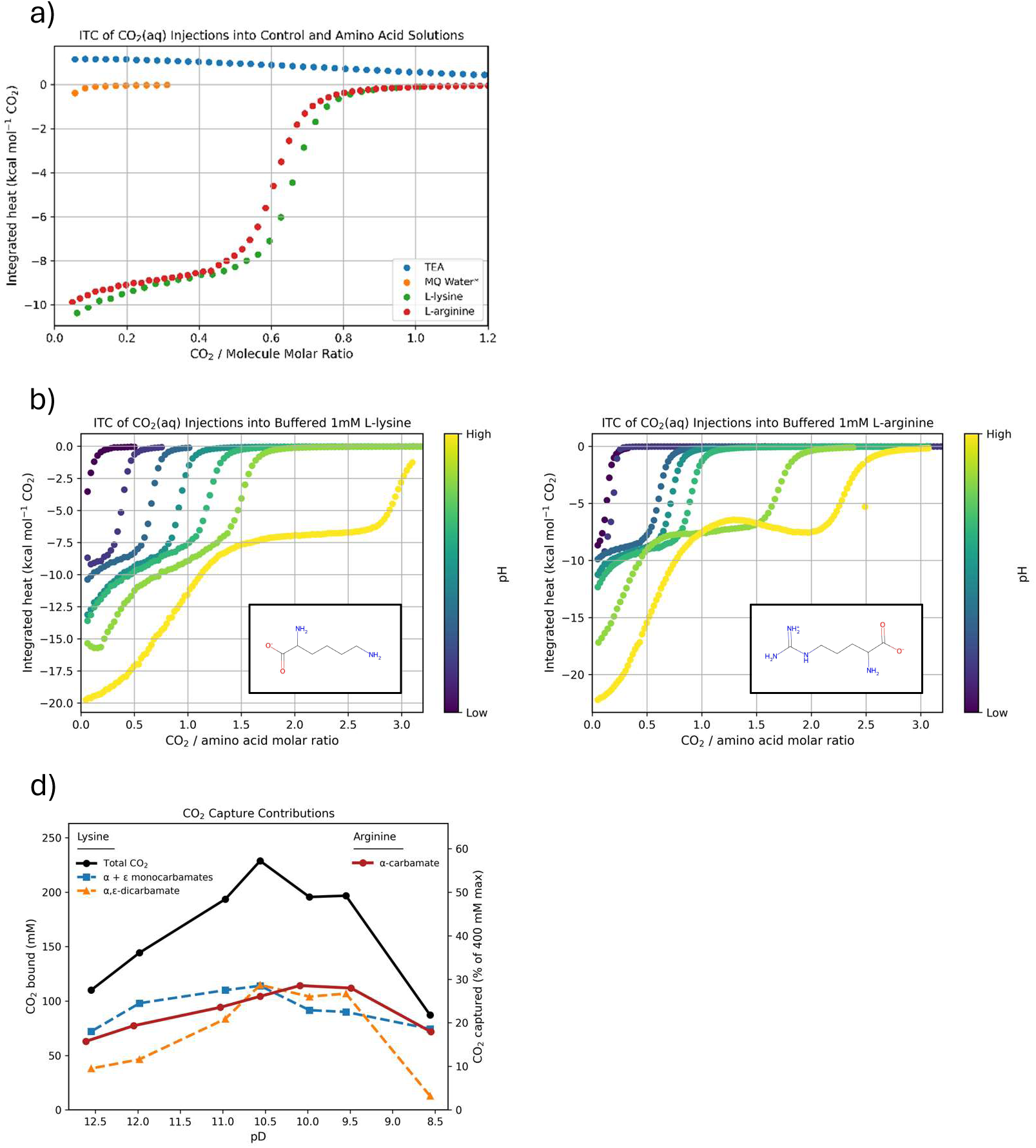
- Calorimetric signatures and carbamate speciation of CO_₂_ uptake by Lys and Arg. **a,** ITC titration of CO_₂_(aq) into control and amino acid solutions at their intrinsic pH showed biphasic exothermic responses for L-Lys and L-Arg, but only minor heat signals for MQ water and triethylamine (TEA**). b,c,** pH-adjusted ITC titrations of 1 mM L-Lys **(b)** and L-Arg **(c)** showed that increasing initial pH shifted the calorimetric transitions to higher CO_₂_-to-amino-acid molar ratios, affirming that the observed heats arise from pH-dependent coupled equilibria d, NMR-derived carbamate-bound CO_₂_ contributions for Lys and Arg as a function of pD. Lys formed α-carbamate, ε-carbamate, and α,ε-dicarbamate species, with maximal carbamate loading near pD 10.5, whereas Arg formed only detectable α-carbamate.

To further investigate the origin of this behavior, the calorimetric response of Lys and Arg was measured across an increasing alkalinity range using NaOH and HCl to modulate amine deprotonation and promote carbamate formation (**Figure 2b-c**). As expected, increasing the pH extended the apparent CO_2_-to-amino-acid saturation molar ratio, defined as the ratio at which the calorimetric signal returned to baseline and further CO_2_ injections produced negligible heat. However, the ratio value approaches 3.0, which exceeds the expected stoichiometric limit of 0.5 and 1.0 for monoamine and diamine systems^22^. Classical carbamate formation requires two amines per captured CO_2_ molecule: one forming the carbamate while a second acts as a proton acceptor^22^. For diamine systems such as Lys, the theoretical upper limit becomes 1.0 if both amines act independently. Interestingly, Arg followed a nearly identical trend despite the substantially higher pKₐ of the guanidinium group (∼12.5–13.8)^23^ compared to the Lys ε-NH₃⁺ group (∼10.5), which should significantly limit Arg’s CO_2_ capture potential per amine in the pH region tested as the guanidinium will be mostly protonated. Additional ITC experiments were performed on N-α-acetyl Lys and Arg at their intrinsic pH, pH 6.4 and 6.8 respectively, and compared with their parent amino acids (**Figure S3**). The negligible calorimetric response following α-amine acetylation indicates that the enthalpic signal in the parent amino acids cannot be explained by side-chain chemistry alone and requires α-amine participation together with carbonate chemistry. In tandem, control titrations of CO_2_ into NaOH (**Figure S4**) produced qualitatively similar biphasic profiles, although with a reduced second transition that became pronounced only at higher NaOH concentration. These observations indicate that biphasic calorimetric behavior is not unique to amino acid systems but reflects intrinsic features of CO_2_ reactivity in basic aqueous solutions.

To directly identify the CO_2_-derived species underlying this behavior and quantify their distribution, we therefore turned to NMR spectroscopy.

### NMR Reveals Dual-Site Carbamate Formation and Enhanced CO_2_ Capture

^1^H and ^13^C NMR spectroscopy were performed at higher Lys concentration than the ITC experiments (200 mM). Here CO_2_ gas was bubbled into a NaOH-buffered L-Lys solution (pD 13.32) in D_2_O and aliquoted at different pD values. The samples were then measured with both ^1^H NMR and ^13^C NMR.

Under very basic condition at pD 12.55, ^13^C NMR spectra revealed that CO_2_ was converted to carbonate/bicarbonate at 168.4 ppm and L-Lys carbamates around 164 – 165 ppm (**Figure S5)**. The signal of carbonate/bicarbonate was observed to shift gradually to 161 ppm when the pD was decreased to 8.56, indicating the pH-dependent fractions of them as reported previously in a ^13^C NMR titration study^24^. Interestingly, rather than carbonate/bicarbonate, the carbamates were detected as major products at pD 12.55 consisting of α-, ε-, and α, ε-dicarbamate, probably because the carbamate formation is about two orders of magnitude faster than CO_2_ hydration^25,26^. The formation of the dicarbamate species peaked around pD 10.56. Thanks to the high field ^1^H NMR spectra, it was possible to quantify the fractions of the three carbamate species at different pDs **(Figure S6)**. pD ≈ 10.56 represents a critical point at which the total carbamate population reaches a maximum. Below this pD, carbamate formation is dominated by the α-amino group, whereas above pD 10.56 the ε-carbamate becomes the predominant species. The populations of both α- and ε-carbamates decrease sharply as the solution pH approaches their respective pKₐ values (with pH ≈ pD − 0.4)^27^, consistent with protonation-driven destabilization of the carbamate species^20,28,29^. Notably, even at high pD values (> pKa of ε-NH_2_) where substantial deprotonation of the α-amino group is expected, the ε-carbamate formation outcompeted the α-amino group, which we attribute to either a higher kinetic rate or higher carbamate equilibrium constant (*K*_carb_).

In terms of carbon capture, the NMR-derived speciation data were converted from carbamate concentrations into carbamate-bound CO_2_ concentrations by accounting for the stoichiometry of each species, with monocarbamates corresponding to 1 mol CO_2_ per mol species and dicarbamates corresponding to 2 mol CO_2_ per mol species. The resulting CO_2_ loading was reported both as an absolute concentration of carbamate-bound CO_2_ (mM) and as a capacity-normalized value, defined as the fraction of the theoretical maximum carbamate capacity assuming full conversion of all available amine sites (400 mM maximum) (**Figure 2c, Figure S6**). This normalization allows direct comparison of carbamate-derived CO_2_ loading across pD values. The total carbamate-bound CO_2_ reaches a maximum of 57.2% of the theoretical amine-site capacity, corresponding to 228.8 mM CO_2_. This increased loading is associated with enhanced formation of the dicarbamate species, which increases the amount of CO_2_ bound per amino-acid molecule.

To assess whether this behavior is general, analogous NMR experiments were performed on CO_2_-treated L-Arg under similar alkaline conditions (**Figure 2d**). In contrast to Lys, no evidence for side-chain carbamate formation was detected. Instead, the spectra revealed the presence of bicarbonate species together with α-carbamates, suggesting that carbamate formation in Arg is restricted to the α-amino group.

ITC, pH titration, and NMR provide complementary information on the CO_2_ capture process: ITC measures the overall heat of reaction, pH titrations constrain the underlying acid–base equilibria, and NMR validates the predicted reaction speciation. However, these techniques cannot independently deconvolute the overlapping thermodynamic contributions of the individual reaction steps. To achieve this, we develop a mechanistic model describing each reaction involved in CO_2_ capture.

### Thermodynamic Modelling Accurately Describes ITC Transitions

To interpret the ITC heat signals in terms of CO_2_-induced speciation, we developed a thermodynamic equilibrium model describing CO_2_ injection into neutral to basic aqueous amino acid solutions (**Figure S7**). The model couples CO_2_ hydration, carbonate speciation, amine protonation, and carbamate formation for Lys and Arg, together with their associated reaction enthalpies. Rather than treating α,ε-dicarbamate formation as a separate reaction, it is represented by an effective cooperativity constant (K_double_), allowing dicarbamate formation to emerge from sequential occupation of the α- and ε-amino groups (**Figure S8**). Following each CO_2_ injection, the coupled equilibria are solved subject to mass and charge balance to determine solution speciation, reaction heats, and pH, enabling direct comparison with experimental ITC thermograms, pH titrations, and NMR-derived speciation (**Figure S8**). The model also resolves the otherwise inaccessible enthalpic contributions from protonation, carbamate formation, carbonate speciation, and cooperative dicarbamate formation.

Model parameters were determined by globally fitting ITC datasets collected between pH 8 and 11 for Lys, Arg, and NaOH controls using a hierarchical weighted least-squares procedure. Initial pH values were optimized individually to account for uncertainty in starting proton activity, while parameter confidence intervals were assessed by profile likelihood analysis which estimates confidence bounds by varying each parameter while reoptimizing all others. Carbonate pathway enthalpies were constrained using NaOH controls (**Figures S9–S10**), and ε-site protonation parameters were constrained using N-α-acetyl-L-Lys (**Figure S11**) before fitting the remaining parameters. Model performance was then evaluated by comparing simulated and experimental pH titrations across multiple initial pH conditions.

Overall, the fitted simulations reproduced the major features and trajectories of both the ITC thermograms and pH titrations for Lys (**Figures S12-15)** and Arg (**Figures S16-18)** across the investigated alkalinity range, supporting that the simulator captures the dominant processes governing CO_2_ uptake (**Figure 3a-b**). Importantly, agreement between the heat and pH data constrains the fitted parameters beyond enthalpy alone and reduces the likelihood of obtaining non-unique solutions that reproduce thermograms but fail to describe solution speciation (**Figure 3c** and **Figure S19).** Deviations between simulation and experiment were most pronounced at the early injection regime of higher pH datasets. These discrepancies likely arise from processes not explicitly represented in the current model, particularly the contribution of base-catalyzed CO_2_ hydration, which becomes increasingly important under strongly alkaline conditions^30^. Hydroxide-mediated CO_2_ hydration may accelerate the conversion of dissolved CO_2_ into bicarbonate under strongly alkaline conditions, leading to an underestimation of the initial heat response and deviations in the predicted speciation^31^. Despite these local deviations, the model accurately captures the transitions in the thermograms described earlier as well as showcasing that the second sigmoidal transition is a result of buffer collapse, not carbamate binding saturation (**Figure 3d**). This is evident from the overlaid pH profile showing a sharp pH decrease coinciding with the second thermal transition. Taken together, these results indicate that the reaction network is sufficient to explain the experimental ITC and pH behavior, while remaining consistent with the independently observed NMR speciation. This provides a mechanistic framework for interpreting the individual thermodynamic contributions to CO_2_ capture.

**Figure 3.**
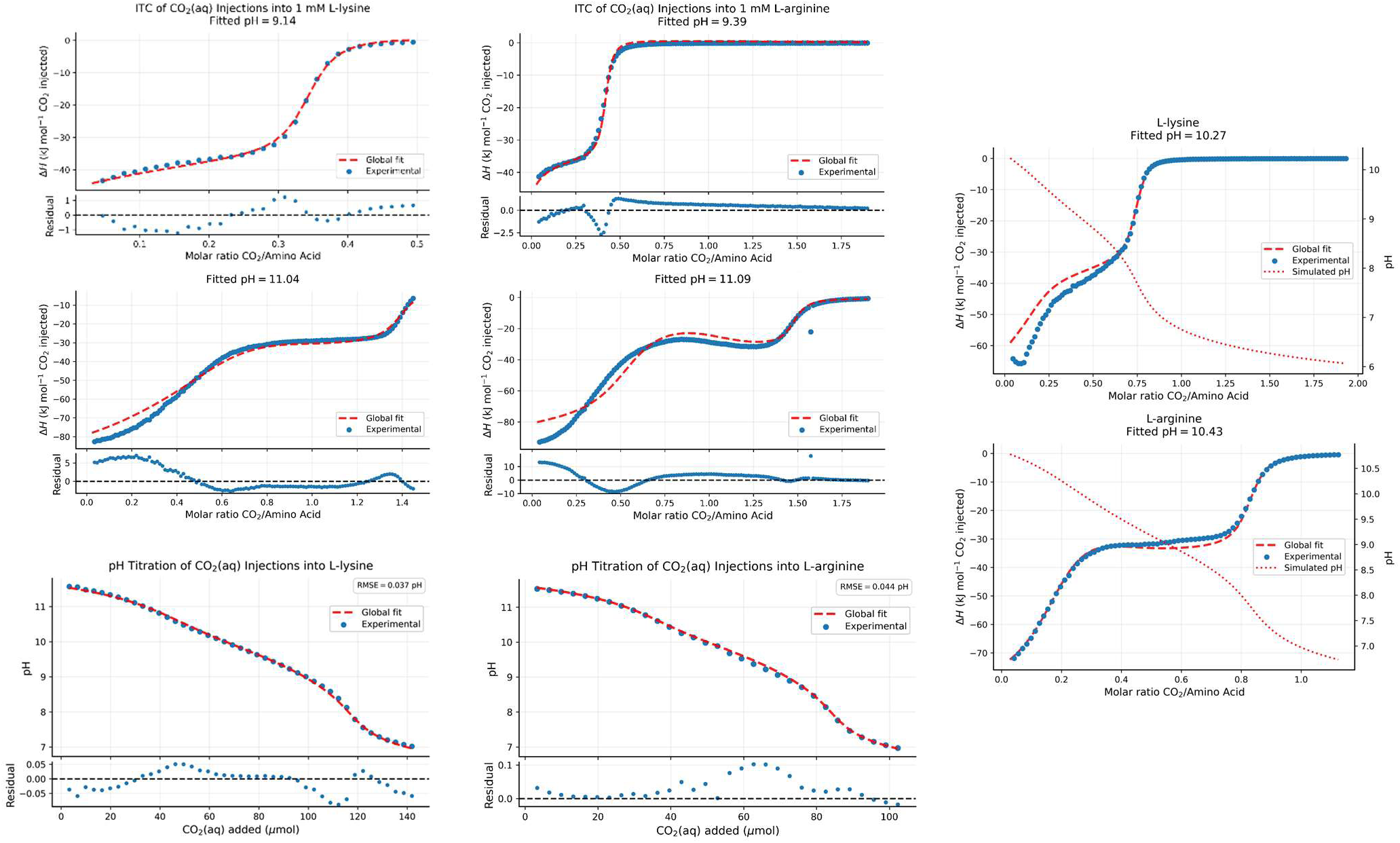
- Thermodynamic model reproduces CO_₂_-induced heat and pH transitions in Lys and Arg. **a,b,** Global equilibrium-model fits to representative ITC thermograms for 1 mM L-Lys **(a)** and L-Arg **(b)** at low and high initial pH, with residuals shown below each fit**. c,** The same fitted parameter set was used to simulate pH titration profiles for Lys and Arg during progressive CO_₂_ addition. Agreement between the calorimetric and pH datasets constrains the model beyond the heat signal alone and supports that the dominant coupled equilibria are captured. **d,** Overlay of simulated pH profiles with fitted ITC thermograms shows that the second sigmoidal heat transition coincides with a sharp pH decrease, consistent with loss of buffering capacity rather than carbamate-binding saturation.

### Carbamate Stability Is Governed by Site-Specific Free Energies in Lys and Arg

**Table 1 (Table S2)** summarizes the fitted thermodynamic parameters and their derived Δ*G*^∘^and Δ*S*^∘^values for the CO_2_ system and for protonation and carbamate formation in Lys and Arg. The carbonate equilibria were fixed at logK = 6.4 for HCO_3−_ and logK = 10.3 for CO3²^−^, with fitted protonation enthalpies (ΔH°) of −9.4 and −14.2 kJ mol−1, respectively, matching literature values of -8.95 and -15.4 kJ mol^-1^ within the margin of error ^13^.

**Table 1.**
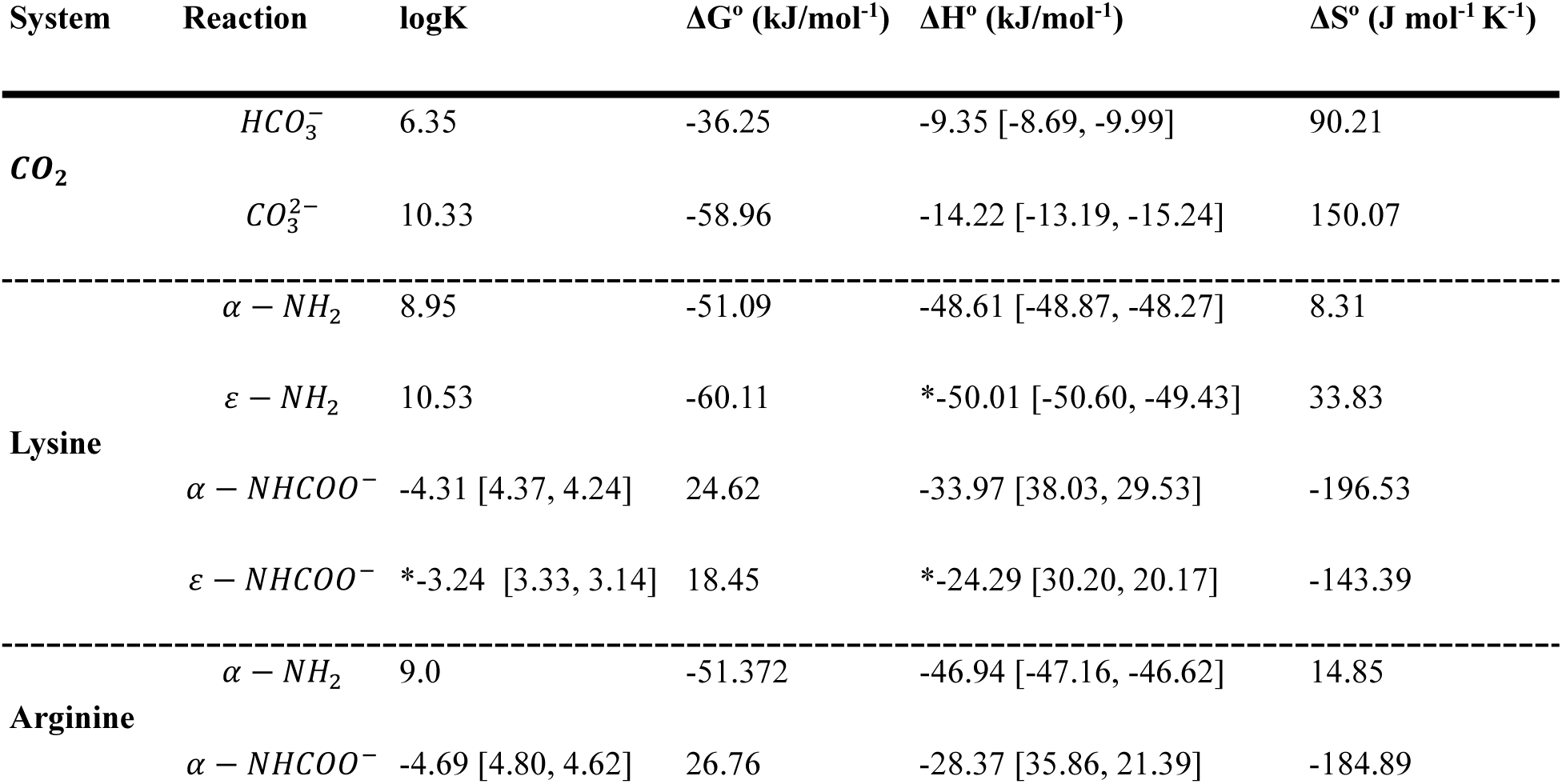
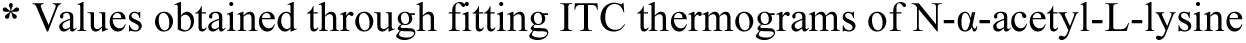
- Fitted thermodynamic parameters for carbonate protonation, amino-group protonation, and carbamate formation equilibria. Values are reported for the reactions written in the direction shown in the reaction column. Carbonate equilibria are written in the protonation direction, HCO₃⁻ + H⁺ ⇌ CO_2_(aq) + H₂O and CO₃²⁻ + H⁺ ⇌ HCO₃⁻. Amino-group protonation constants refer to R–NH₂ + H⁺ ⇌ R–NH₃⁺, while carbamate constants refer to carbamate formation from the neutral amine, R–NH₂ + CO_2_(aq) ⇌ R–NHCOO⁻ + H⁺. Thus, for carbamate formation, larger logK values, i.e. less negative values in this table, correspond to more favorable carbamate formation. Bracketed values indicate confidence intervals from profile likelihood analysis. Asterisks denote values obtained from fits to N-α-acetyl-L-lysine. No values were obtained for the guanidinium on Arg.

| System | Reaction | logK | $\Delta G^\circ$ (kJ/mol<sup>-1</sup>) | $\Delta H^\circ$ (kJ/mol<sup>-1</sup>) | $\Delta S^\circ$ (J mol<sup>-1</sup> K<sup>-1</sup>) |
| --- | --- | --- | --- | --- | --- |
| **CO<sub>2</sub>** | $HCO_3^-$ | 6.35 | -36.25 | -9.35 [-8.69, -9.99] | 90.21 |
| | $CO_3^{2-}$ | 10.33 | -58.96 | -14.22 [-13.19, -15.24] | 150.07 |
| **Lysine** | $\alpha - NH_2$ | 8.95 | -51.09 | -48.61 [-48.87, -48.27] | 8.31 |
| | $\varepsilon - NH_2$ | 10.53 | -60.11 | *-50.01 [-50.60, -49.43] | 33.83 |
| | $\alpha - NHCOO^-$ | -4.31 [4.37, 4.24] | 24.62 | -33.97 [38.03, 29.53] | -196.53 |
| | $\varepsilon - NHCOO^-$ | *-3.24 [3.33, 3.14] | 18.45 | *-24.29 [30.20, 20.17] | -143.39 |
| **Arginine** | $\alpha - NH_2$ | 9.0 | -51.372 | -46.94 [-47.16, -46.62] | 14.85 |
| | $\alpha - NHCOO^-$ | -4.69 [4.80, 4.62] | 26.76 | -28.37 [35.86, 21.39] | -184.89 |

\* Values obtained through fitting ITC thermograms of N-α-acetyl-L-lysine

The fitted thermodynamics for Lys and Arg show that protonation enthalpies are comparatively similar across the amino groups, whereas carbamate formation is much more site dependent. For Lys, protonation of the α- and ε-amino groups is nearly the same (Δ*H*^∘^ = −49 and −50 kJ mol^-1^, respectively), while Arg α-protonation is only slightly less exothermic ( −47 kJ mol^-1^). The distinction is more pronounced in the carbamate-forming equilibria: Lys α-carbamate formation is more exothermic than ε-carbamate formation (Δ*H*^∘^ = −34 versus −25 kJ mol^-1^), but it incurs a much larger entropy penalty (Δ*S*^∘^ = −200 J mol^-1^ K^-1^versus −140 J mol^-1^ K^-1^). As a result, the ε-carbamate is thermodynamically less costly overall on a standard-state basis (Δ*G*^∘^ = 18 kJ mol^-1^) than the α-carbamate (25 kJ mol^-1^). However, K_double_ was not identifiable because of strong correlation with the α- and ε-carbamate equilibria. As a result, the reported α- and ε-carbamate thermodynamics should be interpreted as effective parameters that may partially incorporate stabilization from α,ε-dicarbamate formation.

The ε-carbamate was not identifiable for Arg, and only α-carbamate formation thermodynamics could be fitted, consistent with the absence of an NMR-detectable side-chain carbamate. This is most likely due to the guanidinium’s highly elevated pK_a_ of ∼12.4, well above even the highest pH reached during our titration experiments. The thermodynamics of Arg α-carbamate formation followed nearly the same trend as Lys but is less favorable (**Table 1**), additionally matching the trend in NMR with a higher max α-carbamate population in Lys (**Figure S6b**).

Although the fitted standard free energies for α- and ε-carbamate formation are positive, they describe the isolated carbamate-forming equilibrium as defined in **Fig S7** and therefore do not represent the overall driving force for CO_2_ capture in solution. Carbamate accumulation is instead shaped by the surrounding equilibrium network, including amine protonation, proton transfer, hydroxide neutralization, CO_2_ hydration, and bicarbonate/carbonate speciation. In this sense, the apparent formation of carbamate is coupled to the ability of the solution to accept and redistribute protons, as in conventional zwitterionic descriptions of amine–CO_2_ chemistry^17^. Thus, carbamate concentration should not be interpreted as an intrinsic property of the amine site alone, but as an outcome of both the carbamate-forming equilibrium and the broader solution environment. However, when the surrounding equilibria are held constant, differences in the fitted carbamate-forming free energies provide a meaningful measure of the relative propensity of each amine site to form carbamate. A site with a smaller free-energy penalty, or equivalently a larger logK_carb_ under the present definition, is therefore expected to give a higher carbamate population under otherwise comparable conditions.

The Lys and Arg carbamate constants fall within the same broad stability regime as other amines, but with clear site-specific differences^13,20^. Here, greater carbamate stability corresponds to larger (i.e., less negative) values of logK_carb_, since the equilibrium constant is defined in the direction of carbamate formation (**Fig S7**). Literature logK_carb_ span approximately from −6.0 to −3.7 at 298 K, with most amines clustering between about −5.6 and −3.6. The Lys α-carbamate (log *K*_carb_ = −4.3) is comparable to the MEA/PIPD/PIPZ, whereas the Arg *α*-carbamate (−4.7) is slightly less stable and sits closer to the cyclic and alkanolamine carbamates. In contrast, the Lys ε-carbamate (log *K*_carb_ = −3.2) lies beyond the upper end of the literature values, suggesting that the Lys side-chain amine forms particularly stable carbamate species relative to the α-amino group and performs comparably to the most favorable carbamate-forming amines reported in literature. This trend agrees with previous studies where carbamate stability primarily attributed to steric accessibility and solvent reorganization rather than amine basicity^20,32^. Consistent with this interpretation, the fitted parameters show that the larger exothermicity of carbamate formation does not necessarily correspond to greater carbamate stability. Although Lys α-carbamate formation is more exothermic than ε-carbamate formation, it is also associated with a substantially larger unfavorable entropy contribution. Consequently, the α-carbamate has a less favorable standard free energy than the ε-carbamate, despite its more exothermic formation enthalpy. The higher fitted stability of the ε-carbamate is therefore best described by its lower overall free-energy penalty rather than by the magnitude of the heat released during formation.

### Thermodynamic Modeling Identifies Optimal CO_2_ Capture Conditions

Having established that the fitted thermodynamic model accurately reproduces the experimental CO_2_ capture behavior, we next used it as a predictive tool to map the experimental design space and identify conditions that maximize CO_2_ retention. Rather than experimentally testing every combination of operating variables, the validated model enables rapid in silico evaluation of how CO_2_ capture responds to changes in temperature, initial pH, Lys concentration, and CO_2_ loading. The optimized Lys parameter set (**Table 1, Table S2**) was therefore used to generate two-dimensional heat maps predicting CO_2_ retention across these variables (**Figure 4**), providing a framework for identifying optimal operating conditions and distinguishing whether capture is dominated by carbonate chemistry, amine-mediated retention, or carbamate formation. Temperature effects were approximated through a van’t Hoff treatment (**Figure S20)**^33^ of the fitted equilibria, allowing the extracted enthalpies to influence simulated capture behavior. For each condition, three factors were examined: total CO_2_ retention, retention achievable by NaOH alone, and the resulting amine-mediated retention calculated as the difference between total and NaOH capture (Figure 20). Additionally, carbamate-specific heat maps were produced to isolate the contribution of carbamate formation from amine-mediated capture.

**Figure 4.**
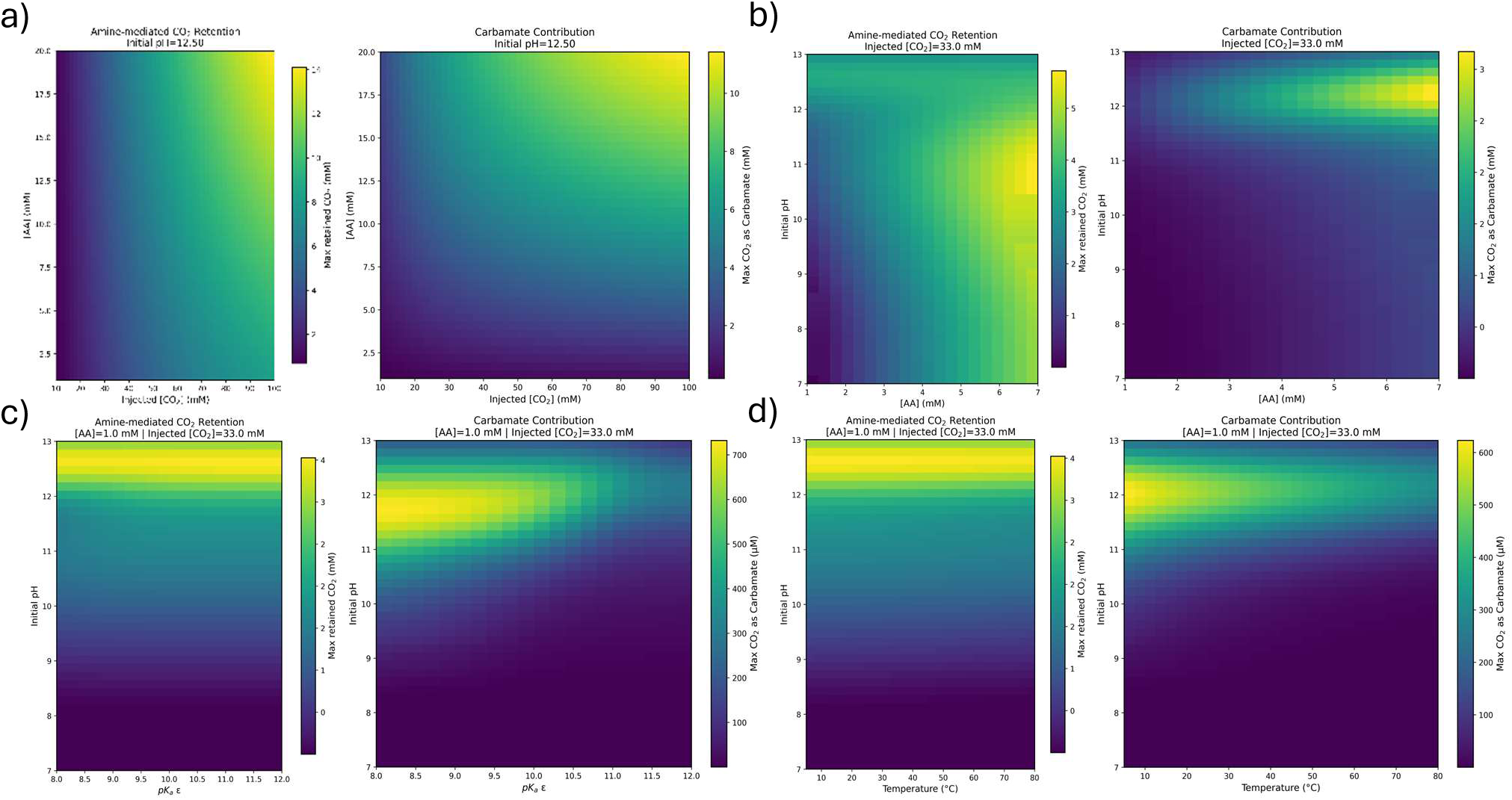
- Simulated Lys design-space sweeps for amine-mediated CO_2_ retention and carbamate formation. Heat maps generated from the optimized Lys parameter set. Each panel pair shows a two-parameter sweep, where the variables shown on the axes were varied across the indicated ranges while the remaining conditions were held fixed. In each panel, the left heat map shows amine-mediated CO_2_ retention, defined as total retention minus the NaOH-only background, while the right heat map shows CO_2_ retained as α-, ε-, or α,ε-carbamate species. **a**, Supplied CO_2_ and Lys concentration were swept at fixed initial pH 12.50. **b**, Initial pH and Lys concentration were swept at fixed supplied CO_2_ of 33.0 mM. **c**, Initial pH and ε-amine pKₐ were swept at fixed Lys concentration of 1.0 mM and supplied CO_2_ of 33.0 mM. **d**, Initial pH and temperature were swept at fixed Lys concentration of 1.0 mM and supplied CO_2_ of 33.0 mM. Across the sweeps, carbamate formation occupies a narrower optimum than amine-mediated retention and is favored by sufficient Lys concentration, alkaline pH, lower ε-amine pKₐ, and lower temperature.

Across all parameter sweeps, total CO_2_ retention peaked at high pH and showed behavior closely resembling the NaOH-only simulations (**Figure S21a)**. Capture increased sharply above ∼ pH 11 and reached its highest value at the highest pH tested (pH 13), indicating that most total retained carbon originates from hydroxide-driven carbonate and bicarbonate formation rather than direct amine binding. This trend was nearly the same for temperature and pK_a,ε_, (**Figure S21b-c)** consistent with carbonate equilibria contributing most strongly to the total capture capacity. While capture is significant in this pH region, achieving these strongly alkaline solutions is costly and unfeasible in industrial settings. This can be mitigated by increasing the amount of CO_2_ delivered to the system. At the intrinsic pH of Lys (pH ∼9.4) the total captured CO_2_ is about 20x lower than at pH 12.5.

Subtracting the NaOH background revealed a different landscape for amine-mediated capture, which is defined here as additional capture capacity of the solution when amine is present. This includes amine-mediated proton shuffling between the amines and the carbonate pathway as well as carbamate formation. Amine-mediated CO_2_ retention remained relatively small in magnitude, roughly 4 to 5 times less retained CO_2_ than total retention **(Figure 4a-d left)**, except in the case of high amine and high CO_2_ loading which showed about half as much (**Figure 4a left, Figure S21d)**. This is indication that CO_2_ retention is limited by amine concentrations; high amounts of amine and CO_2_ supply allow for the amine-mediated capture to begin competing with NaOH-mediated capture.

The strongest amine contribution occurred within a narrower operating window centered around pH 11 and increasing Lys loading rather than continuously increasing with pH (**Figure 4b left)**. This behavior reflects the competing requirements for carbamate formation. For CO_2_ capture via carbamates, sufficient amine deprotonation is required to enable nucleophilic attack on CO_2_, but excessively alkaline conditions increasingly divert dissolved CO_2_ toward carbonate formation and reduce the relative contribution from amine chemistry. This is partially mitigated by increasing amine concentration and supplying more CO_2_.

The carbamate contribution maps further emphasized this distinction. Unlike total retention, carbamate contribution to capture showed a much narrower band of optima **(Figure 4b-d right)**. Lower temperatures consistently increased carbamate populations **(Figure 4d right)**, in agreement with the fitted negative carbamate formation enthalpies for α- and ε-carbamate formation indicating thermodynamically favored carbamate stabilization at reduced temperature through the van’t Hoff relationship. Carbamate formation was also maximized at a pH higher and lower pK_a,ε_ than amine-mediated retention and showed a clear dependence on Lys concentration **(Figure 4b-c right)**, consistent with the requirement for available deprotonated amine sites. Additionally, the retention amount for carbamate capture is again around 4-5 times lower than amine-mediated retention, except for in the same case of increased amine and CO_2_ loading **(Figure 4a right)**. This suggests that CO_2_ capture in aqueous amino acid systems is dominated primarily by carbonate and bicarbonate speciation, with the amines acting mainly as a base. Under the investigated conditions, carbamate therefore represents a secondary capture pathway rather than the dominant storage form and only becomes a major contributor when sufficient CO_2_ loading is achieved. For example, at an amino acid concentration of 10 mM, approximately 40 mM supplied CO_2_ is required before carbamate-bound CO_2_ accounts for 50% of the retained CO_2_. At lower amino acid concentrations, this threshold is reached only at increasingly higher CO_2_ loadings.

The separation between total and amine-mediated capture further suggests that maximizing total CO_2_ uptake and maximizing carbamate utilization are fundamentally different optimization problems. In practice, this distinction may explain why many aqueous amine systems achieve high overall capture despite comparatively weak carbamate equilibria^20,32,34,35^.

### Peptide Engineering of Carbamate Formation

Although NMR revealed that the α-carbamate is the predominant carbamate species formed by L-lysine, interest in CO_2_ capture by proteins motivated a closer examination of lysine side-chain reactivity, as lysine ε-amino groups are generally the most solvent-exposed and readily accessible amines in proteins. Because carbamate stabilization is expected to depend on the presence of a nearby second amine, the spacing between lysine side chains may be an important determinant of carbamate formation and stability. To extend the amino-acid framework to a peptide context, a series of N-terminally acetylated Lys- and Arg-based model peptides was designed with the general sequence ACE-GK(G)_n_KG and ACE-GR(G)_n_RG, where *n* denotes the number of intervening glycine residues. Increasing the number of glycine residues systematically increased both the average Lys–Lys separation from 5.64 to 10.07 Å and the corresponding amine–amine distance from 8.90 to 12.61 Å, providing a controlled system for evaluating the effect of inter-amine spacing on CO_2_ capture (**Table 2**).

**Table 2.**
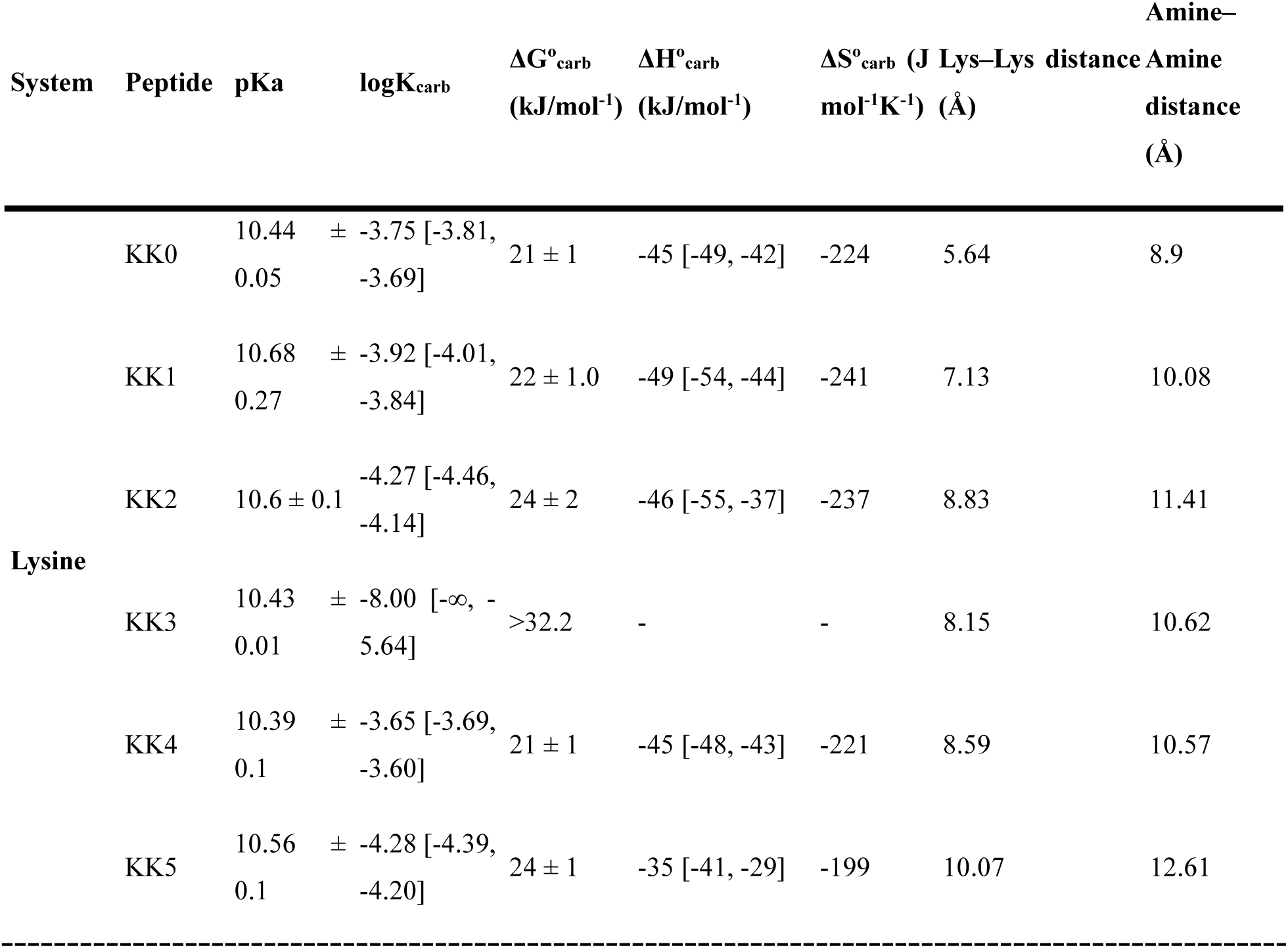
- Thermodynamic parameters and structural descriptors for the Lys-containing peptide series. The reported pKₐ values are Boltzmann-weighted averages predicted from CREST conformational ensembles using PROPKA and correspond to the amine deprotonation equilibrium R–NH₃⁺ ⇌ R–NH₂ + H⁺. Lys–Lys and amine–amine distances are Boltzmann-weighted average distances obtained from the same conformational ensembles. The reported logKcarb, ΔG°carb, ΔH°carb, and ΔS°carb describe the apparent carbamate-forming equilibrium, R–NH₂ + CO_2_(aq) ⇌ R–NHCOO⁻ + H⁺, using a shared carbamate-forming equilibrium for the two Lys side chains within each peptide. Larger logKcarb values (i.e., less negative values) correspond to more favorable carbamate formation. Bracketed values indicate 95% confidence intervals obtained from profile likelihood analysis. Em dashes indicate parameters that were not identifiable from the fitted thermograms because the corresponding confidence intervals were effectively unbounded.

ITC measurements were collected for each peptide and analyzed using the thermodynamic framework developed for the amino acid systems (**Figures S22–S23**). Initial amine pKₐ values were estimated from CREST-generated conformational ensembles followed by Boltzmann-weighted PROPKA calculations. Across the Lys peptide series, the predicted pKₐ values remained similar (10.35–10.73), preventing independent resolution of the two ε-amino groups during fitting. Consequently, the model employed a shared apparent pKₐ, carbamate equilibrium constant, and carbamate formation enthalpy for the two lysine side chains within each peptide.

The fitted thermodynamic parameters are summarized in Table 2. Relative to free Lys, all peptide constructs exhibited less favorable carbamate formation, with fitted logK_carb_ values ranging from −3.65 to −4.27, while carbamate formation could not be reliably identified for KK3. However, carbamate stability showed no systematic dependence on either Lys–Lys separation or amine–amine distance. For example, KK2 exhibited one of the least favorable carbamate equilibria (logK_carb_ = −4.27), whereas KK4 partially recovered carbamate stability (−3.65) despite having a larger average separation between lysine residues. Similarly, KK5, with the largest average spacing, did not exhibit the weakest carbamate equilibrium. These results indicate that varying lysine spacing alone is insufficient to predict or systematically improve carbamate stability within this peptide series.

Comparison with the amino acid results further illustrates the influence of the peptide environment. The fitted carbamate equilibrium for free Lys ε-carbamate (logK_carb_ = −3.24) was more favorable than all peptide values and comparable to the strongest carbamate-forming amines reported in the literature. In contrast, the peptide carbamate equilibria fell within the broader range reported for aqueous amines, indicating that incorporation of the lysine side chain into a peptide generally reduces the apparent stability of the carbamate-forming equilibrium.

For the Arg peptide series, the guanidinium side chain remained too basic to permit independent determination of carbamate thermodynamics, with fitted pKₐ values reaching the upper limit of the fitting range (>13.4), consistent with the high basicity of arginine. In addition, satisfactory fits required the apparent carbonate and bicarbonate reaction enthalpies to be reoptimized rather than fixed to the NaOH-derived values (**Figures S24-25**). Although the origin of this behavior cannot be uniquely assigned, it indicates that the thermodynamic parameters describing the carbonate pathway may not be directly transferable from simple NaOH solutions to all peptide systems.

Overall, these results demonstrate that the thermodynamic framework remains applicable to peptide systems while showing that inter-amine spacing alone does not determine carbamate stability. Within this series, modifying lysine separation did not provide a systematic strategy for enhancing carbamate formation, suggesting that additional structural features beyond average amine spacing will need to be considered when designing peptide- or protein-based CO_2_ capture systems.

## Discussion

### Isothermal Titration Calorimetry as a Quantitative Probe of CO_2_ Capture Thermodynamics

This work establishes an experimentally constrained thermodynamic framework for understanding CO_2_ interactions with amino acids and peptides in aqueous environments while demonstrating the broader utility of isothermal titration calorimetry (ITC) as a quantitative tool for studying aqueous CO_2_ capture. More broadly, the results demonstrate that ITC, when combined with thermodynamic modelling and independent pH titration and NMR measurements, can monitor how the coupled CO_2_–amine equilibrium network evolves during progressive CO_2_ loading, including contributions from hydration, proton transfer, carbonate speciation, and carbamate formation.

Although ITC directly measures only the net heat released following each CO_2_ injection, coupling the calorimetric data to an equilibrium model transforms these heat signals into a quantitative description of the underlying chemistry. The agreement between simulated and experimental thermograms, independent pH titrations, and NMR-derived speciation demonstrates that the fitted reaction network captures the dominant thermodynamic processes governing CO_2_ uptake. Rather than interpreting calorimetric measurements solely as reaction enthalpies, the present framework establishes ITC as a method for resolving the evolution of solution speciation during CO_2_ loading while simultaneously extracting reaction-specific thermodynamic parameters that cannot be measured directly.

This capability extends beyond the amino acid systems examined here. Because the methodology relies only on calorimetric measurements together with an appropriate thermodynamic description of the solution chemistry, it provides a general strategy for quantitatively analyzing other aqueous amine-based capture systems and determining how individual chemical processes contribute to the overall capture behavior.

### Thermodynamic Interpretation Reveals Design Principles for Aqueous CO_2_ Capture

A central outcome of this work is that apparent CO_2_ binding behavior and total carbon retention should not be interpreted directly as measures of carbamate formation. Although carbamates were directly observed and quantified by NMR, both the thermodynamic analysis and simulated capture landscapes showed that carbon storage in aqueous amino acid systems is dominated by carbonate and bicarbonate speciation, with amines primarily regulating proton activity and directing carbon partitioning through the surrounding equilibrium network. This interpretation agrees with previous studies of aqueous amine capture, which have similarly emphasized that carbamate formation competes with, and is thermodynamically coupled to, surrounding acid-base and hydration equilibria rather than functioning as an isolated binding process^13,20,34,35^.

Rather than identifying amino acids as inherently superior CO_2_ sorbents, the present results suggest an alternative design principle in which the surrounding molecular environment governs capture behavior as much as intrinsic amine affinity. Within the aqueous equilibrium regime examined in this work, CO_2_ uptake was controlled primarily by proton redistribution and carbon partitioning between carbamate and inorganic carbon species rather than by carbamate stability alone. In practical terms, this means optimizing not only the intrinsic carbamate stability of an amine site, but also its apparent pKₐ, local charge environment, hydration structure, and ability to participate in proton transfer. The most effective capture scaffolds are therefore unlikely to be those that simply maximize carbamate affinity, but rather those that position amine pKₐ values within the relevant operating pH window while providing neighboring proton-accepting groups or solvent environments that stabilize either carbamate or bicarbonate formation depending on the desired balance between capture capacity and regeneration energy.

This perspective also explains several observations made throughout the present study. Lys exhibited more favorable carbamate formation than Arg despite similar protonation thermodynamics, while the peptide series demonstrated that simply increasing the spacing between lysine residues did not systematically improve carbamate stability. Together, these results indicate that optimizing aqueous CO_2_ capture requires engineering the complete thermodynamic environment surrounding reactive amines rather than modifying a single molecular property. Peptides and proteins provide an attractive route toward this type of optimization because sequence design can independently tune amine spacing, local charge density, side-chain identity, hydration, and proton-transfer pathways while maintaining an aqueous capture medium.

### Experimentally Constrained Models Enable Predictive CO_2_ Capture Design

Beyond interpreting the present experiments, the developed equilibrium model provides a predictive platform for analyzing CO_2_ capture under conditions that are difficult or impractical to access experimentally. By integrating calorimetry, pH evolution, and chemical speciation into a unified framework, the model enables extraction of otherwise inaccessible thermodynamic contributions and prediction of capture behavior across environmental and molecular variables.

The resulting parameter set allows multidimensional operating spaces to be explored computationally using experimentally constrained thermodynamics. Rather than using the model simply to reproduce measured thermograms, the present work demonstrates how it can be used to identify operating conditions that maximize specific capture objectives, including total CO_2_ retention, amine-mediated capture, carbamate formation, or the balance between capture and regeneration. Such experimentally validated simulations provide a practical route for narrowing experimental design spaces before laboratory testing and therefore offer a complementary strategy to conventional solvent screening approaches.

From an engineering perspective, the framework therefore suggests a workflow in which candidate amine systems are not ranked solely according to total CO_2_ uptake, but according to how carbon partitions between carbamate and bicarbonate/carbonate species under realistic operating conditions. Candidate molecules can first be modelled, their thermodynamic parameters determined from ITC measurements, and their capture behavior subsequently predicted across a broad range of operating conditions before experimental optimization. In this way, ITC becomes not only a characterization technique but also a means of parameterizing predictive models capable of guiding the rational development of aqueous CO_2_ capture systems.

### Toward Rational Biomolecular Carbon Capture

The successful extension of the framework from free amino acids to peptide systems further demonstrates that equilibrium-based descriptions remain informative as molecular complexity increases, even when individual reaction pathways become increasingly difficult to isolate experimentally. Although the peptide constructs investigated here did not outperform the corresponding free amino acids, they illustrate how the methodology can be applied to increasingly complex molecular architectures and provide experimentally derived thermodynamic constraints for systems in which direct interpretation of calorimetric data would otherwise be difficult.

Future extensions of this framework could move beyond equilibrium descriptions by incorporating structural information, molecular dynamics, and reaction kinetics into the thermodynamic model. Combined with advances in computational peptide and protein engineering, such approaches could enable rational design of biomolecular architectures that control CO_2_ uptake through engineered local environments rather than optimization of intrinsic binding affinity alone. The present framework therefore provides not only a mechanistic description of aqueous CO_2_–amine chemistry, but also an experimental and computational platform for developing next-generation biomolecular carbon capture systems.

## Acknowledgements

The authors thank Albert Thor Thorhallsson for his support and Dr. Kasper Enemark-Rasmussen for the NMR measurements at the NMR center of Technical University of Denmark. This work is generously supported by the Novo Nordisk Foundation CO2 Research Center (CORC) (project application no. 2022-0031).

## Materials and Methods

### Isothermal Titration Calorimetry

ITC experiments were performed using a MicroCal VP-ITC instrument at 25°C. Amino-acid solutions were prepared at 1 mM in degassed Milli-Q water and adjusted to the desired initial pH using NaOH or HCl. Control experiments were prepared in an identical manner.

Aqueous CO_2_ solutions were prepared by sparging degassed Milli-Q water with pure CO_2_ gas for 10 min at a flow rate of 5 mL/min to achieve near-saturation at RT (∼33 mM). Prior to use, the saturated CO_2_ solution was diluted 2:1 (v/v) with degassed Milli-Q water to obtain the final titrant solution.

The sample cell was loaded with amino-acid solution or control, while the reference cell contained degassed Milli-Q water. The diluted CO_2_ solution was loaded into the injection syringe. Successive injections of 1.5-2.0 μL were performed until no further heat changes were observed and the calorimetric signal returned to baseline. A reference power of 33 μcal s⁻¹ and a stirring speed of 220 rpm was used throughout all experiments.

### pH Titrations

pH titrations were performed at room temperature in 30 mL amino-acid or control solutions prepared in degassed Milli-Q water. Amino-acid solutions were prepared at 1 mM and adjusted to the desired initial pH using NaOH or HCl. Control solutions containing only Milli-Q water were prepared in an identical manner.

The titrant consisted of degassed Milli-Q water continuously sparged with pure CO_2_ throughout the experiment to maintain a constant dissolved CO_2_ concentration (∼33 mM). Titrations were performed in a beaker covered with parafilm, and pH was monitored using a benchtop pH meter. Successive 100 μL injections of the CO_2_ solution were added every 30 s while stirring. The pH was recorded immediately before each injection.

Titrations were continued until the pH approached the bicarbonate buffering region and further CO_2_ additions produced only minimal changes in pH.

### NMR

Liquid ^1^H and ^13^C NMR spectra were measured with an 18.8 T Bruker AVANCEIII HD spectrometer (n_1H_ = 800.18 MHz and n_13C_ = 201.17 MHz) equipped with a 5 mm TCI CryoProbe (Bruker). For the ^1^H spectra, a p/6 excitation pulse was employed with an interscan delay of 7 seconds. For the 13C spectra, a p/6 excitation pulse was employed with an interscan delay of 12 seconds and inverse-gated decoupling. All measurements were conducted at 298 K and chemical shifts are reported relative to TMS (d = 0.0 ppm) using the lock signal as secondary reference. A solution of L-Lys (200 mM, 20 mL) in D₂O was slowly bubbled with CO_2_ (10 vol% CO_2_ / 90 vol% N₂) at a flow rate of 80 mL min⁻¹, while the pD was continuously monitored using a pH meter at room temperature. Aliquots (0.5 mL) were withdrawn at different pD values for NMR analysis. The fractions of the three carbamate species were determined by integrating the corresponding α-or ε-proton resonances and normalizing each to the sum of the respective proton integrals.

### Peptide Design and Procurement

Peptides were designed with the following sequences: ACE-GK(G)_n_KG for lysine-based peptides, and ACE-GR(G)_n_RG for arginine-based peptides. ACE corresponds to an acetylated N-terminus, and the n corresponds to the number of glycines between the lysines or arginines. All peptides were commercially synthesized by GenScript and supplied at ≥95% purity, except for KK2, which was obtained at 76% purity due to synthesis limitations.

### Peptide Conformational Sampling and pKₐ Prediction

The Lys- and Arg-containing peptide series, ACE-GK(G)_n_KG and ACE-GR(G)_n_RG, were constructed to systematically vary the spacing between reactive side-chain amines by increasing the number of intervening glycine residues. Initial peptide geometries were generated and subjected to conformational sampling using CREST (Conformer–Rotamer Ensemble Sampling Tool) v3.0^36^ employing the GFN2-xTB semiempirical Hamiltonian. Conformers within 25 kJ mol⁻¹ of the global minimum were retained for subsequent analysis.

For each peptide, the lowest-energy conformers were analyzed using PROPKA v3.5.1^37^ to predict the pKₐ values of the lysine and arginine side-chain amines. Because multiple conformations contribute to the equilibrium structure in solution, pKₐ values were averaged over the conformational ensemble using Boltzmann weighting,

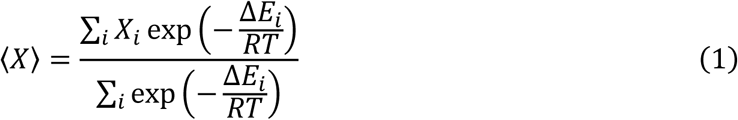

where Δ*E*_i_is the relative energy of conformer *i*, *R*is the gas constant, and *T*is 298.15 K, and where *X* corresponds to either the PROPKA-predicted pKₐ value or the Lys–Lys and amine–amine distances calculated for each conformer. Standard deviations reported in Table 2 represent the Boltzmann-weighted variability across the conformational ensemble.

The same Boltzmann-weighting procedure was applied to calculate the average Lys–Lys and amine–amine distances for each peptide. The Lys–Lys distance was defined as the distance between the geometric centroids of the two lysine residues, whereas the amine–amine distance was defined as the distance between the terminal ε-amino nitrogen atoms (Nζ) of the two lysine side chains. These structural descriptors were used to quantify how increasing the number of intervening glycine residues altered both the overall separation between the lysine residues and the distance between the reactive amine groups responsible for carbamate formation.

The Boltzmann-weighted pKₐ values served as fixed protonation parameters during thermodynamic fitting of the peptide ITC datasets. Because the predicted pKₐ values of the two lysine side chains were similar for all peptides, the calorimetric data could not independently resolve their protonation equilibria. Consequently, each peptide was described using a shared apparent amine pKₐ, a shared carbamate-forming equilibrium constant, and common protonation and carbamate formation enthalpies for the two lysine side chains.

### Thermodynamic model of CO_2_–amine speciation

A thermodynamic equilibrium model was developed to describe CO_2_ uptake in neutral to alkaline aqueous amino acid solutions by coupling carbonate chemistry, amino acid protonation, carbamate formation, and calorimetric heat generation within a unified framework (**Figures S7–S8**). The model was designed to reproduce the evolution of solution speciation during sequential CO_2_ injections while simultaneously predicting calorimetric heats and solution pH. A schematic overview together with the complete mathematical formulation is provided in the Supplementary Information (**Figures S7–S8**).

Carbonate chemistry was described by the coupled equilibria between dissolved CO_2_, bicarbonate, and carbonate together with water autoionization. Amino acid chemistry was represented by the protonation equilibria of each ionizable group and carbamate formation at each reactive amine site. For lysine, independent α- and ε-amine protonation and carbamate formation equilibria were included together with carboxyl deprotonation. Rather than introducing α,ε-dicarbamate formation as a separate reaction pathway, cooperative double carbamation was incorporated through an effective cooperativity constant (K_double_), allowing dicarbamate formation to emerge naturally from sequential occupation of the α- and ε-amine sites. Arginine was treated analogously but without an independently fitted side-chain carbamate equilibrium because no ε-carbamate species were detected experimentally.

Instead of solving each chemical species independently, amino acid speciation was described using a molecular partition-function formalism. Each amino acid was represented as an ensemble of discrete microstates corresponding to every accessible combination of protonation and carbamate occupancy (**Figure S8**). Statistical weights for each microstate were calculated directly from the corresponding equilibrium constants and the instantaneous solution composition. Summation over all microstates produced a molecular partition function from which the equilibrium populations of every amino acid species were obtained. This approach ensured that all protonation and carbamate states remained internally consistent while automatically satisfying amino acid mass balance. The partition function further allowed direct calculation of experimentally inaccessible quantities, including total protonated amine populations, individual α- and ε-carbamate populations, cooperative dicarbamate populations, and the net molecular charge of the amino acid.

The amino acid microstates were coupled to carbonate chemistry through shared proton and carbon mass balances. Following each CO_2_ injection, the dissolved CO_2_ concentration was updated to account for injection dilution before simultaneously solving the coupled equilibrium equations subject to total carbon conservation, amino acid mass balance, and global charge neutrality. Ionic strength was recalculated after each iteration, and activity coefficients were updated using the Davies extension of Debye–Hückel theory until self-consistency was achieved. Initial solution compositions corresponding to the experimentally measured starting pH values were generated by including NaOH or HCl as strong electrolytes within the same mass- and charge-balance framework, allowing all subsequent calculations to proceed from experimentally relevant conditions.

To simulate isothermal titration calorimetry experiments, the equilibrium composition was recalculated after every CO_2_ injection, and the extent of each individual chemical process was determined from the change in species populations between successive equilibrium states. The predicted calorimetric heat was then obtained as the sum of the reaction extents multiplied by their corresponding reaction enthalpies, thereby resolving the individual contributions from protonation, carbamate formation, carbonate speciation, and cooperative dicarbamate formation to the total measured heat signal. Because the model simultaneously enforces charge balance and explicitly tracks proton concentration, the same equilibrium calculation also predicts the corresponding pH trajectory for each experiment without introducing additional fitted parameters. Consequently, a single parameter set can be evaluated simultaneously against experimental ITC thermograms, pH titrations, and independently measured NMR speciation, providing stronger mechanistic constraints than fitting calorimetric data alone. Water autoionization was assigned a fixed deprotonation enthalpy of −55.8 kJ mol⁻¹^38^.

### Thermodynamic Parameter Estimation

Thermodynamic parameters were determined by globally fitting ITC thermograms collected across multiple initial pH values using the equilibrium model described above. Raw ITC thermograms were integrated using NITPIC to obtain normalized injection heats, which were converted to kJ mol⁻¹ of injected CO_2_ prior to analysis. For each experiment, the first injection was excluded to minimize mixing artifacts. Residuals were normalized using an empirical scaling function, and a gradually increasing weighting scheme was applied over the first 5% of injections to reduce the influence of early-injection mixing effects.

Model parameters were estimated using nonlinear weighted least-squares optimization. Rather than fitting every parameter simultaneously, a hierarchical fitting strategy was employed in which progressively more complex experimental systems were used to constrain subsets of parameters before introducing additional degrees of freedom. This approach reduced parameter correlation while ensuring that parameters describing simpler chemistries were independently constrained before fitting more complex reaction networks.

The carbonate subsystem was determined first using NaOH control experiments containing no amines. Carbonate equilibrium constants were initially fixed to literature values^13^ while the bicarbonate and carbonate protonation enthalpies were globally fitted across all NaOH ITC datasets. Following identification of the carbonate enthalpies, these values were fixed and the carbonate equilibrium constants were subsequently refined. The resulting parameter set was then validated against independent NaOH pH titration datasets to verify that both calorimetric and pH behavior were simultaneously reproduced.

Following characterization of carbonate chemistry, N-α-acetyl-L-Lys was used to isolate the ε-amino group. Carbonate parameters determined from the NaOH analysis were fixed, allowing the ε-amine carbamate equilibrium constant, carbamate formation enthalpy, and protonation enthalpy to be determined. Because simultaneous optimization of all three parameters resulted in strong parameter correlations, carbamate equilibrium and enthalpy were first optimized together before fixing these values and independently determining the protonation enthalpy.

The free Lys datasets were subsequently analyzed to determine the α-amino group thermodynamics while retaining the ε-amine parameters obtained from N-α-acetyl-L-Lys. Literature pKₐ values were fixed throughout this stage. The α-carbamate equilibrium constant and protonation enthalpy were initially optimized simultaneously, after which these parameters were fixed and the α-carbamate formation enthalpy was determined independently. Once all reaction enthalpies had been established, they were fixed while the corresponding equilibrium constants were reoptimized. The resulting parameter set was finally validated against independent Lys pH titration datasets.

Arginine datasets were analyzed using the same hierarchical strategy, using the Lys parameter set as initial estimates where appropriate. Only α-carbamate formation was considered because neither NMR nor ITC provided evidence for a resolvable side-chain carbamate equilibrium. N-α-acetyl-L-Arg was not investigated because the guanidinium side chain remains protonated throughout the experimentally accessible pH range, preventing independent determination of its protonation thermodynamics.

The peptide series, ACE-GK(G)_n_KG and ACE-GR(G)_n_RG, were fitted using the corresponding amino acid parameters as initial values. Because the α-amino groups are incorporated into the peptide backbone, only the lysine side-chain amines were considered. Initial pKₐ values were fixed to the Boltzmann-weighted PROPKA predictions described above. The two lysine side chains within each peptide exhibited similar predicted pKₐ values and could not be independently resolved from the calorimetric data. Consequently, each peptide was described using a shared apparent amine protonation equilibrium, a shared carbamate-forming equilibrium constant, and common protonation and carbamate formation enthalpies for the two lysine side chains. The shared carbamate equilibrium constant and protonation enthalpy were first optimized simultaneously before being fixed, allowing the carbamate formation enthalpy to be determined independently. This procedure was repeated for each peptide in the series.

For every optimization, the initial pH of each experimental dataset was treated as a nuisance parameter and refined within predefined bounds around the experimentally measured value to account for uncertainty arising from sample preparation, atmospheric CO_2_ contamination, and pH measurement. Dataset-specific pH values were therefore reoptimized for every trial parameter set during global optimization.

### Validation Using pH Titrations

Independent pH titration datasets were not included directly in the ITC objective function but instead served as an external validation of the fitted thermodynamic parameters. Simulated titrations were generated using experimental injection volumes, titrant concentrations, and initial solution compositions. Agreement between simulated and experimental pH trajectories provided an independent assessment of whether the fitted parameter sets accurately reproduced the coupled acid-base chemistry beyond the calorimetric measurements.

### Profile Likelihood Analysis

Parameter identifiability and confidence intervals were assessed using profile likelihood analysis. Each parameter was sequentially fixed over a predefined range while all remaining adjustable parameters were reoptimized using the identical fitting procedure employed during the original optimization. The resulting residual sum of squares (RSS) was recorded as a function of the fixed parameter value. Approximate 95% confidence intervals were determined using a likelihood-ratio threshold of ΔRSS = 3.84, corresponding to profiling of a single parameter while reoptimizing all remaining parameters. Parameters with bounded profiles were considered identifiable, whereas parameters with effectively unbounded confidence intervals were classified as non-identifiable.

## Abbreviations

Arg: L-arginine
CREST: Conformer–Rotamer Ensemble Sampling Tool
D₂O: deuterium oxide
HCl: hydrochloric acid
HMBC: heteronuclear multiple-bond correlation
HSQC: heteronuclear single-quantum coherence
ITC: isothermal titration calorimetry
KKₙ: lysine-based model peptide series ACE-GK(G)ₙKG
Lys: L-lysine
NaOH: sodium hydroxide
NMR: nuclear magnetic resonance
pD: pH measured in deuterium oxide
PROPKA: protein pKₐ prediction program
RMSE: root-mean-square error
RRₙ: arginine-based model peptide series ACE-GR(G)ₙRG
RSS: residual sum of squares
TEA: triethylamine
TMS: tetramethylsilane.

