## Supplementary Figures for "The Thermodynamics of Biomolecular CO_2_ Capture: Disentangling Equilibria in Amino-Acid-based Systems"

### Slide 1
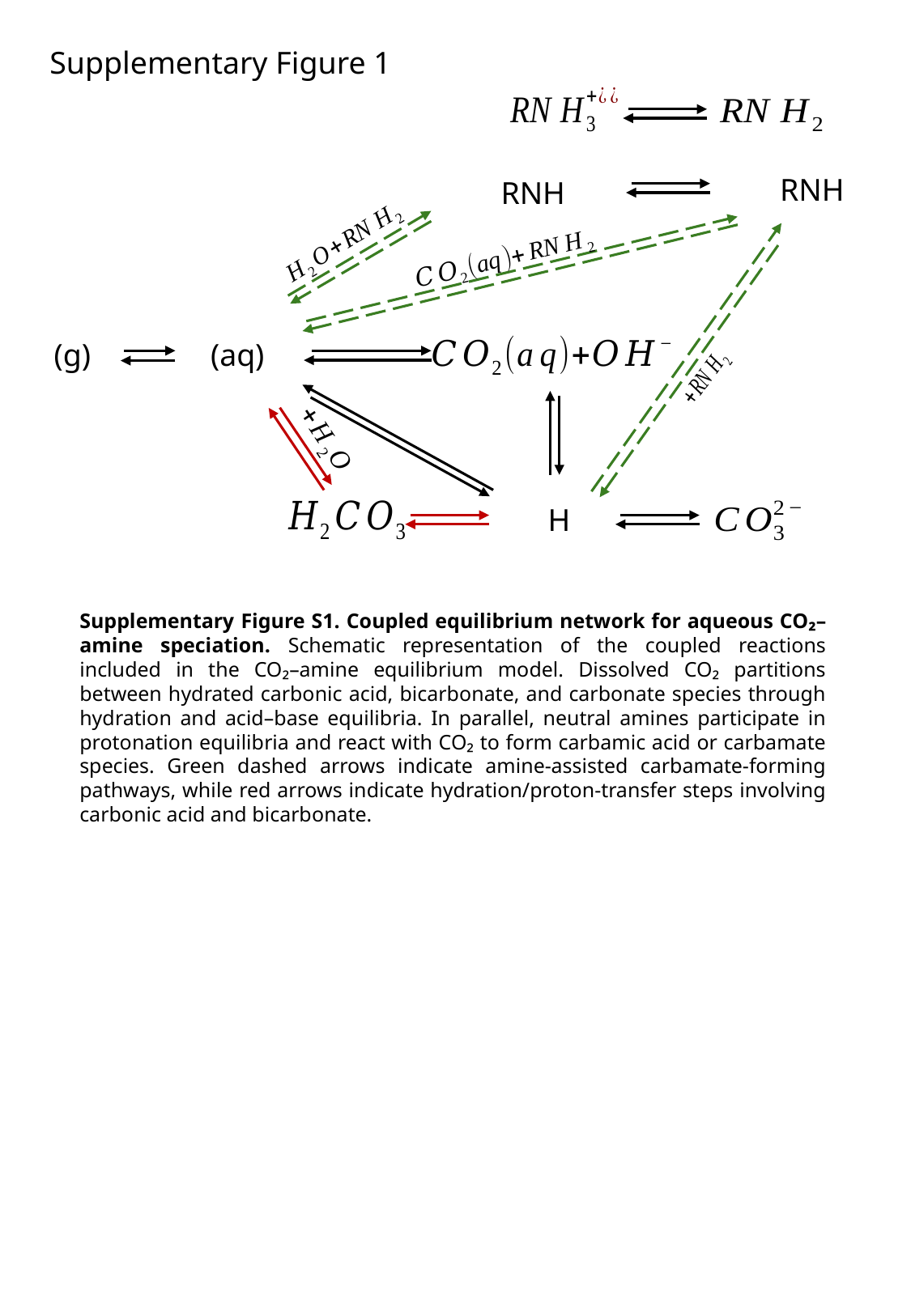

Supplementary Figure 1
Supplementary Figure S1. Coupled equilibrium network for aqueous CO₂–amine speciation. Schematic representation of the coupled reactions included in the CO₂–amine equilibrium model. Dissolved CO₂ partitions between hydrated carbonic acid, bicarbonate, and carbonate species through hydration and acid–base equilibria. In parallel, neutral amines participate in protonation equilibria and react with CO₂ to form carbamic acid or carbamate species. Green dashed arrows indicate amine-assisted carbamate-forming pathways, while red arrows indicate hydration/proton-transfer steps involving carbonic acid and bicarbonate.

### Slide 2
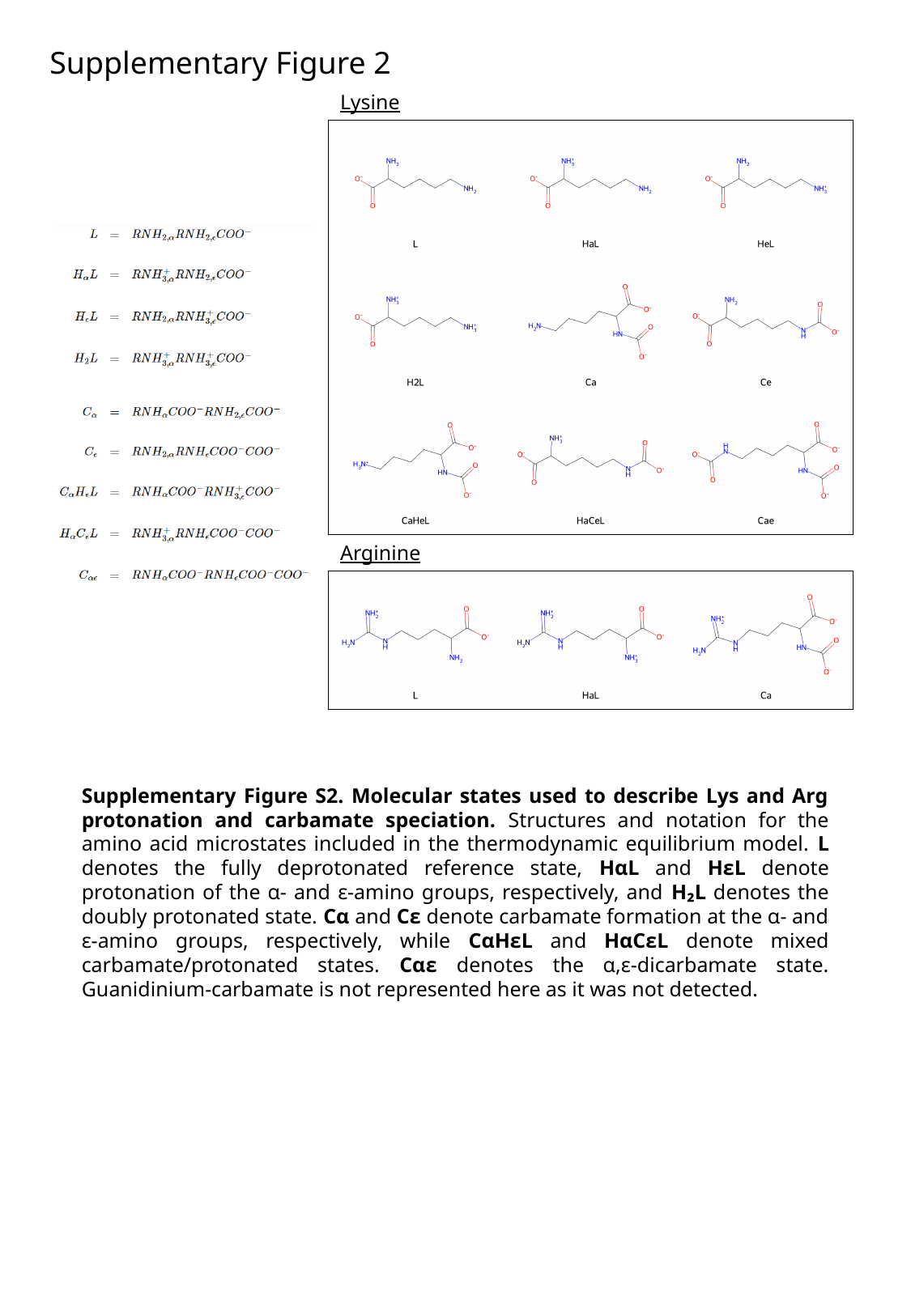

Supplementary Figure 2
Lysine
Arginine
Supplementary Figure S2. Molecular states used to describe Lys and Arg protonation and carbamate speciation. Structures and notation for the amino acid microstates included in the thermodynamic equilibrium model. L denotes the fully deprotonated reference state, HαL and HεL denote protonation of the α- and ε-amino groups, respectively, and H₂L denotes the doubly protonated state. Cα and Cε denote carbamate formation at the α- and ε-amino groups, respectively, while CαHεL and HαCεL denote mixed carbamate/protonated states. Cαε denotes the α,ε-dicarbamate state. Guanidinium-carbamate is not represented here as it was not detected.

### Slide 3
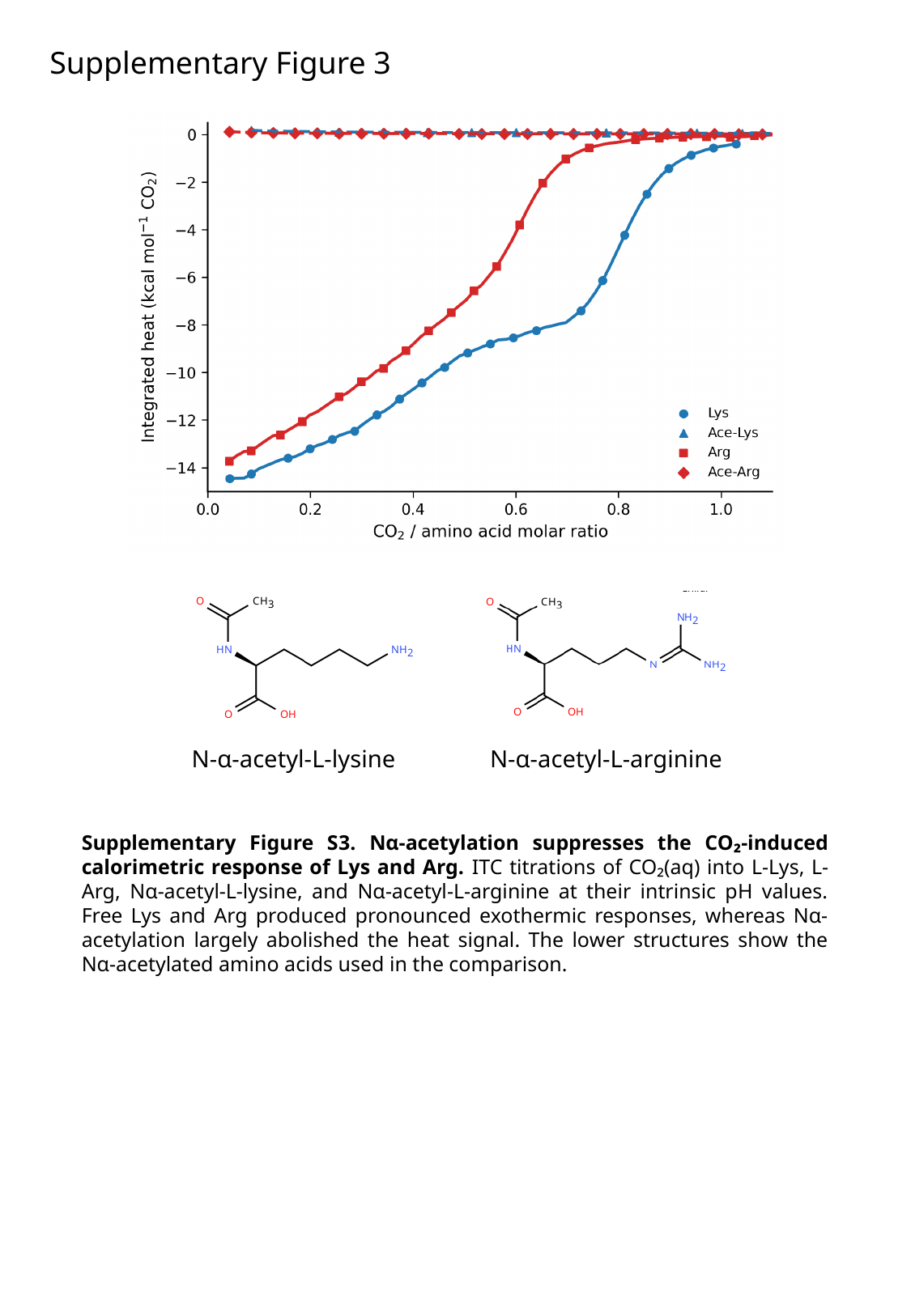

Supplementary Figure 3
N-α-acetyl-L-arginine
N-α-acetyl-L-lysine
Supplementary Figure S3. Nα-acetylation suppresses the CO₂-induced calorimetric response of Lys and Arg. ITC titrations of CO₂(aq) into L-Lys, L-Arg, Nα-acetyl-L-lysine, and Nα-acetyl-L-arginine at their intrinsic pH values. Free Lys and Arg produced pronounced exothermic responses, whereas Nα-acetylation largely abolished the heat signal. The lower structures show the Nα-acetylated amino acids used in the comparison.

### Slide 4
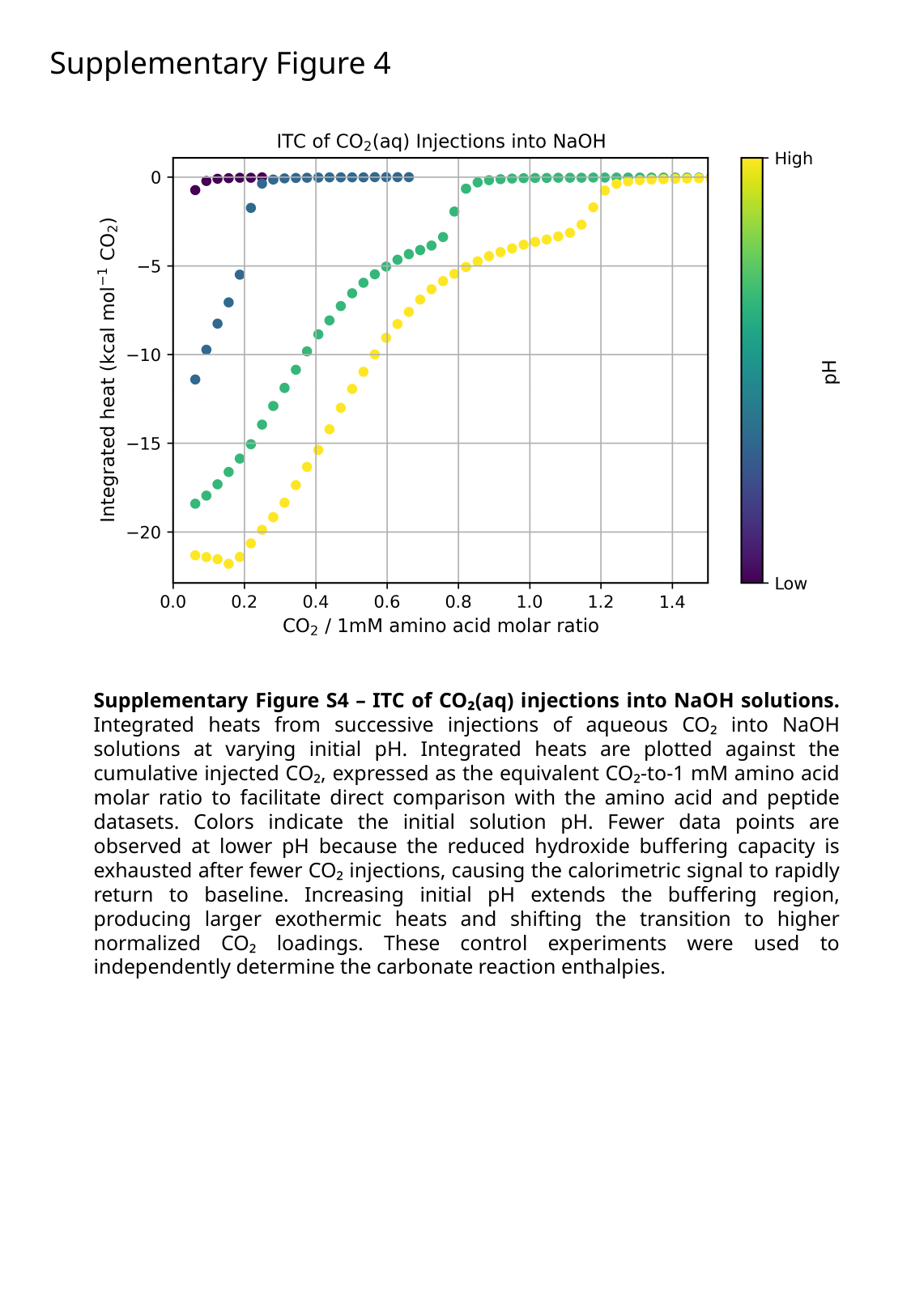

Supplementary Figure 4
Supplementary Figure S4 – ITC of CO₂(aq) injections into NaOH solutions. Integrated heats from successive injections of aqueous CO₂ into NaOH solutions at varying initial pH. Integrated heats are plotted against the cumulative injected CO₂, expressed as the equivalent CO₂-to-1 mM amino acid molar ratio to facilitate direct comparison with the amino acid and peptide datasets. Colors indicate the initial solution pH. Fewer data points are observed at lower pH because the reduced hydroxide buffering capacity is exhausted after fewer CO₂ injections, causing the calorimetric signal to rapidly return to baseline. Increasing initial pH extends the buffering region, producing larger exothermic heats and shifting the transition to higher normalized CO₂ loadings. These control experiments were used to independently determine the carbonate reaction enthalpies.

### Slide 5
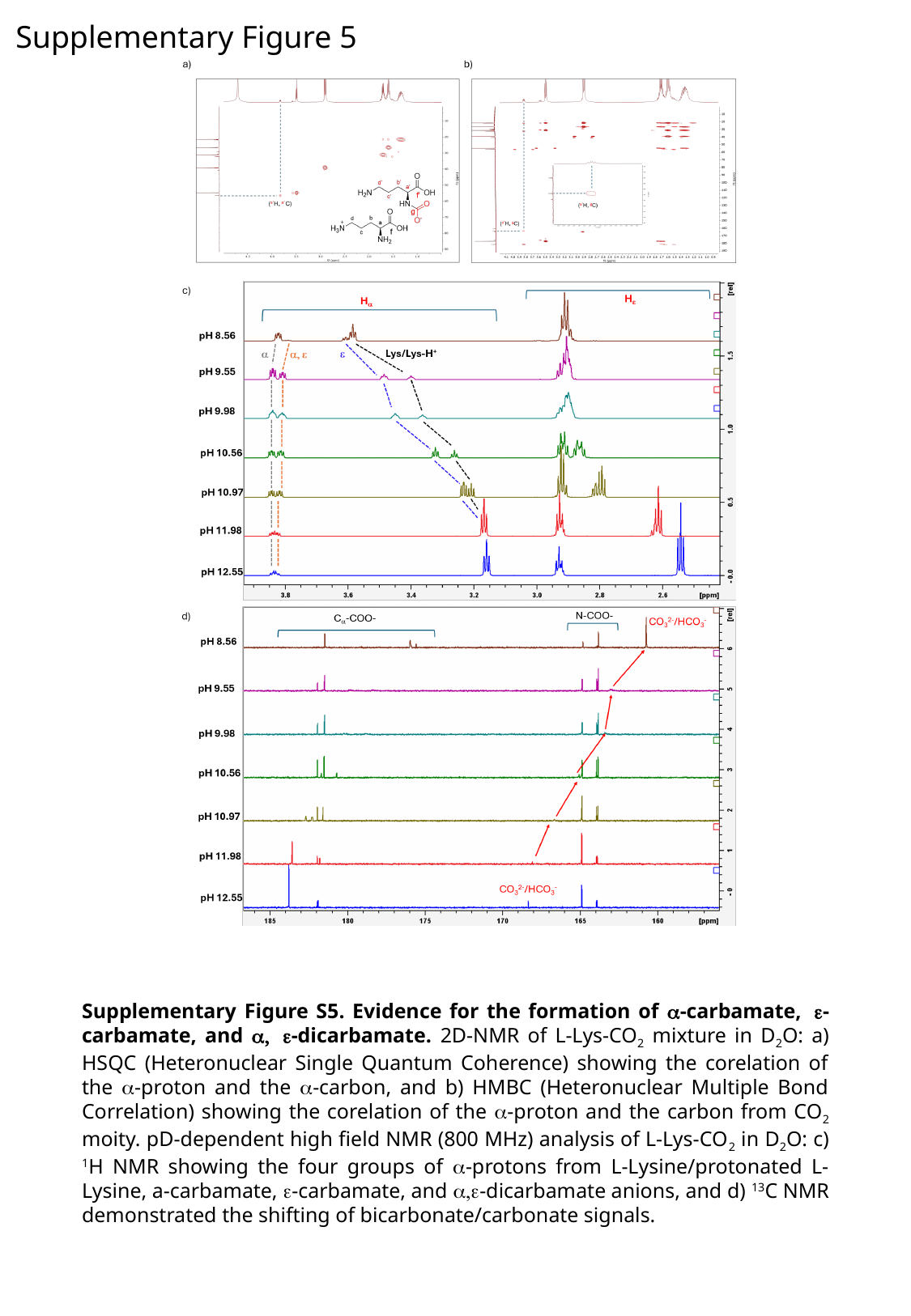

Supplementary Figure 5
Supplementary Figure S5. Evidence for the formation of α-carbamate, ε-carbamate, and α, ε-dicarbamate. 2D-NMR of L-Lys-CO_2_ mixture in D_2_O: a) HSQC (Heteronuclear Single Quantum Coherence) showing the corelation of the α-proton and the α-carbon, and b) HMBC (Heteronuclear Multiple Bond Correlation) showing the corelation of the α-proton and the carbon from CO_2_ moity. pD-dependent high field NMR (800 MHz) analysis of L-Lys-CO_2_ in D_2_O: c) ^1^H NMR showing the four groups of α-protons from L-Lysine/protonated L-Lysine, a-carbamate, ε-carbamate, and α,ε-dicarbamate anions, and d) ^13^C NMR demonstrated the shifting of bicarbonate/carbonate signals.

### Slide 6
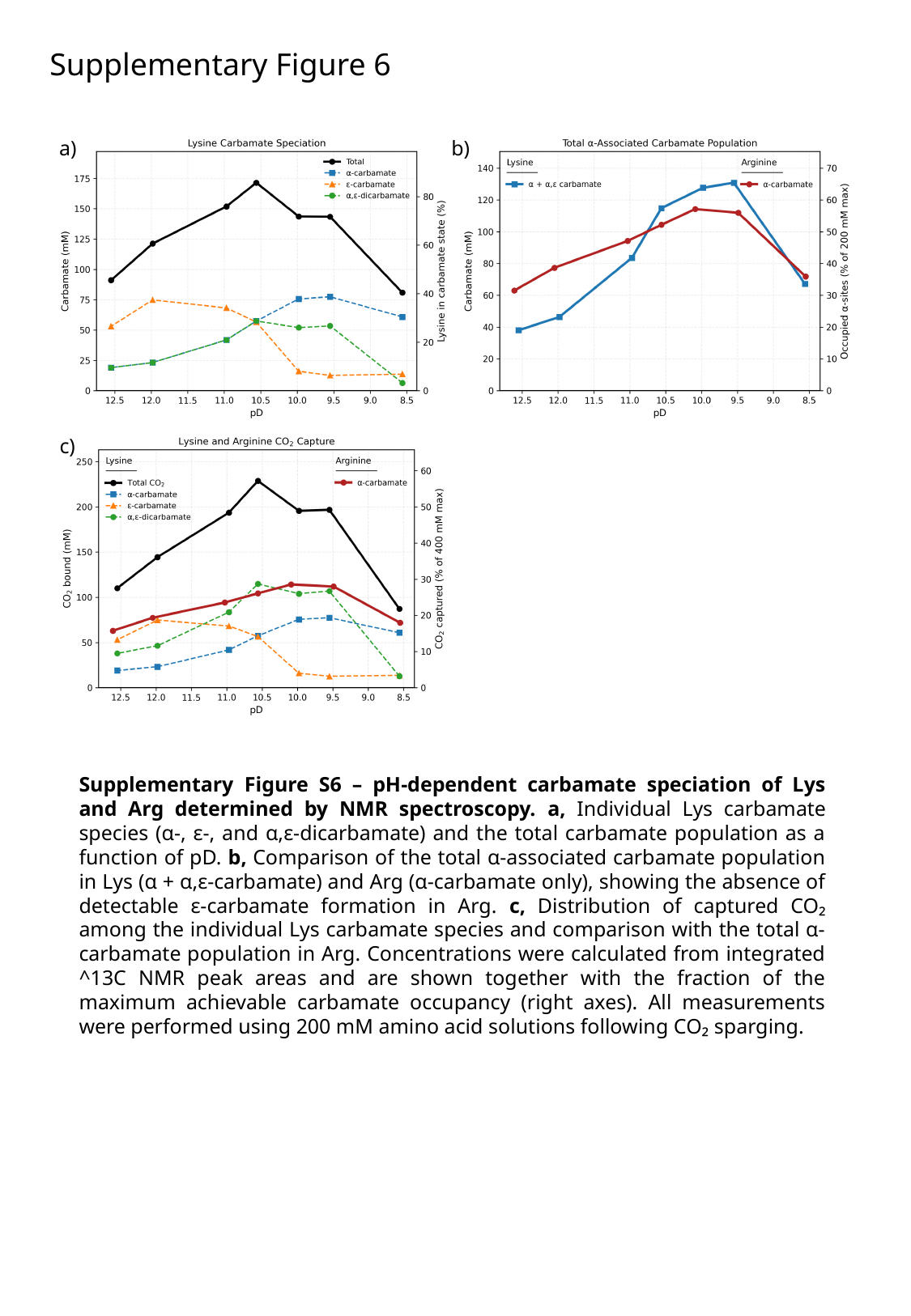

Supplementary Figure 6
b)
a)
c)
Supplementary Figure S6 – pH-dependent carbamate speciation of Lys and Arg determined by NMR spectroscopy. a, Individual Lys carbamate species (α-, ε-, and α,ε-dicarbamate) and the total carbamate population as a function of pD. b, Comparison of the total α-associated carbamate population in Lys (α + α,ε-carbamate) and Arg (α-carbamate only), showing the absence of detectable ε-carbamate formation in Arg. c, Distribution of captured CO₂ among the individual Lys carbamate species and comparison with the total α-carbamate population in Arg. Concentrations were calculated from integrated ^13C NMR peak areas and are shown together with the fraction of the maximum achievable carbamate occupancy (right axes). All measurements were performed using 200 mM amino acid solutions following CO₂ sparging.

### Slide 7
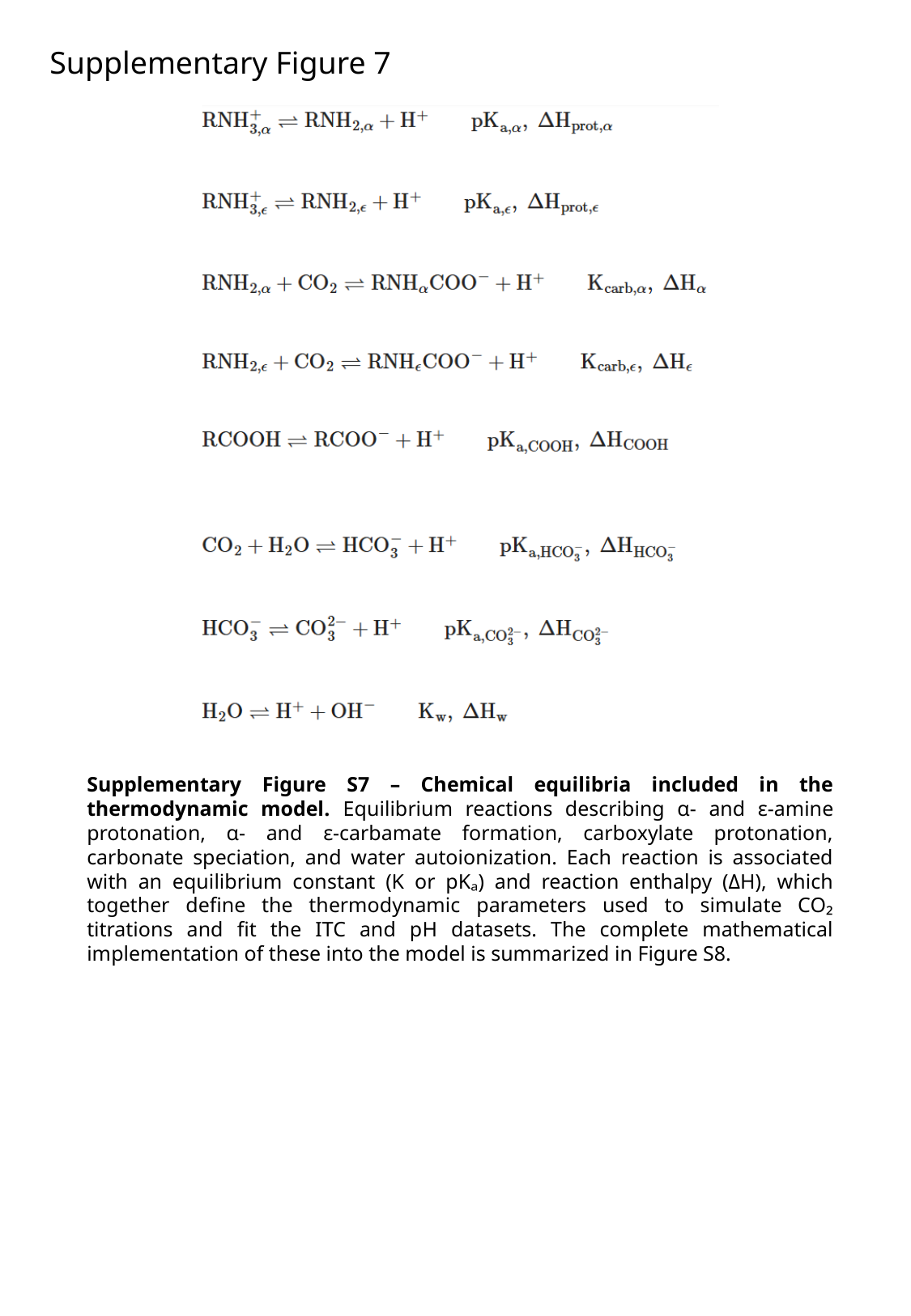

Supplementary Figure 7
Supplementary Figure S7 – Chemical equilibria included in the thermodynamic model. Equilibrium reactions describing α- and ε-amine protonation, α- and ε-carbamate formation, carboxylate protonation, carbonate speciation, and water autoionization. Each reaction is associated with an equilibrium constant (K or pKₐ) and reaction enthalpy (ΔH), which together define the thermodynamic parameters used to simulate CO₂ titrations and fit the ITC and pH datasets. The complete mathematical implementation of these into the model is summarized in Figure S8.

### Slide 8
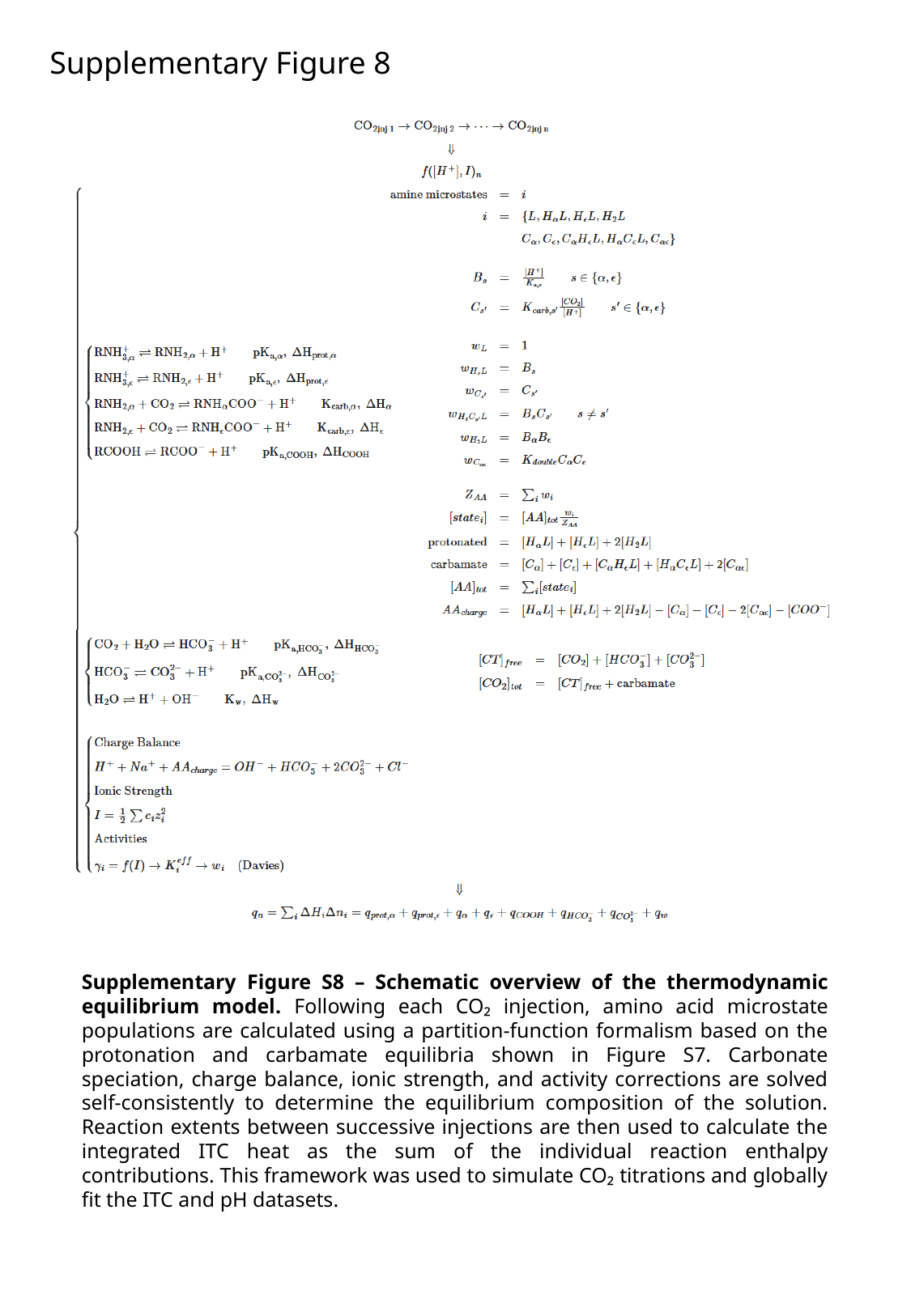

Supplementary Figure 8
Supplementary Figure S8 – Schematic overview of the thermodynamic equilibrium model. Following each CO₂ injection, amino acid microstate populations are calculated using a partition-function formalism based on the protonation and carbamate equilibria shown in Figure S7. Carbonate speciation, charge balance, ionic strength, and activity corrections are solved self-consistently to determine the equilibrium composition of the solution. Reaction extents between successive injections are then used to calculate the integrated ITC heat as the sum of the individual reaction enthalpy contributions. This framework was used to simulate CO₂ titrations and globally fit the ITC and pH datasets.

### Slide 9
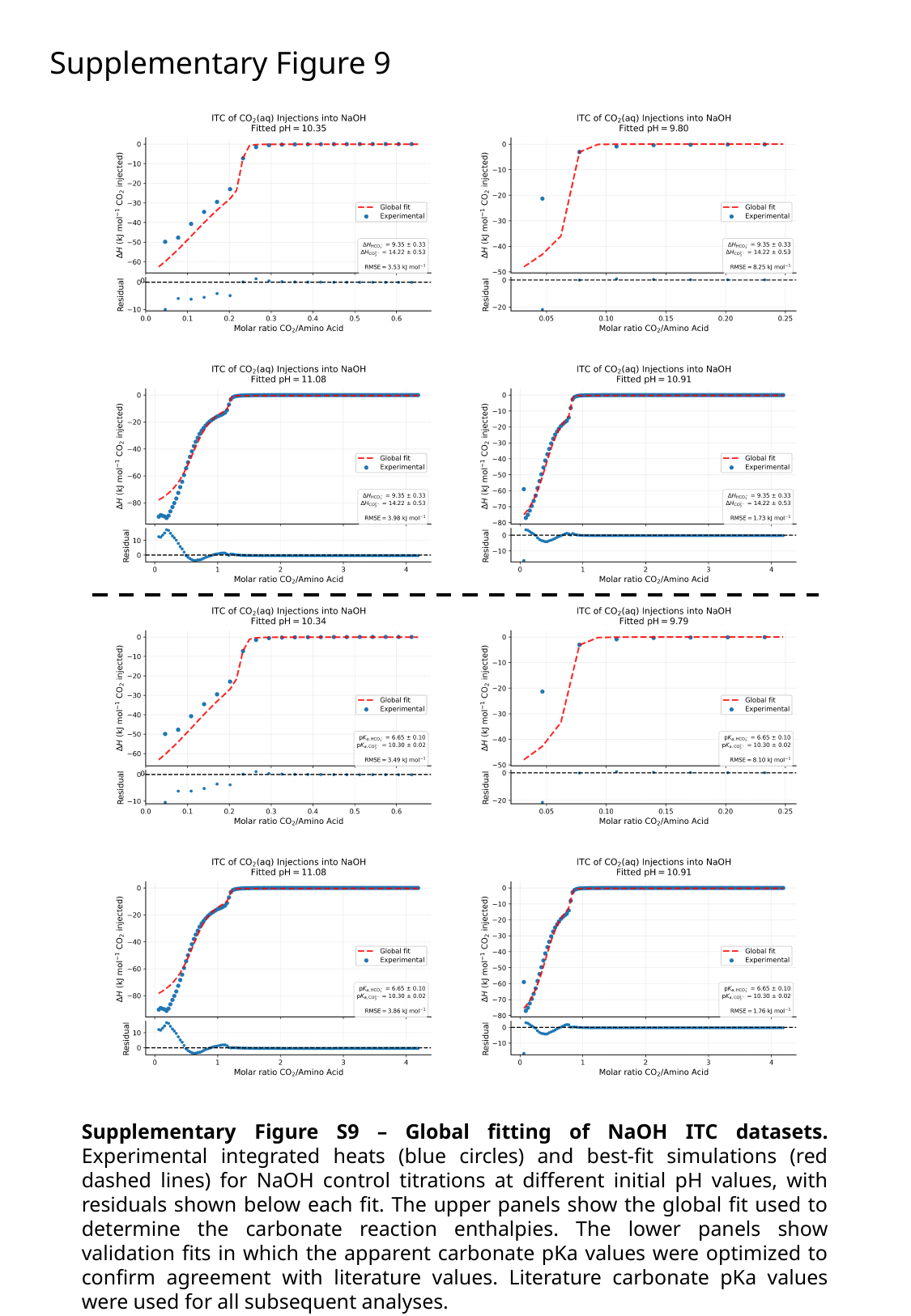

Supplementary Figure 9
Supplementary Figure S9 – Global fitting of NaOH ITC datasets. Experimental integrated heats (blue circles) and best-fit simulations (red dashed lines) for NaOH control titrations at different initial pH values, with residuals shown below each fit. The upper panels show the global fit used to determine the carbonate reaction enthalpies. The lower panels show validation fits in which the apparent carbonate pKa values were optimized to confirm agreement with literature values. Literature carbonate pKa values were used for all subsequent analyses.

### Slide 10
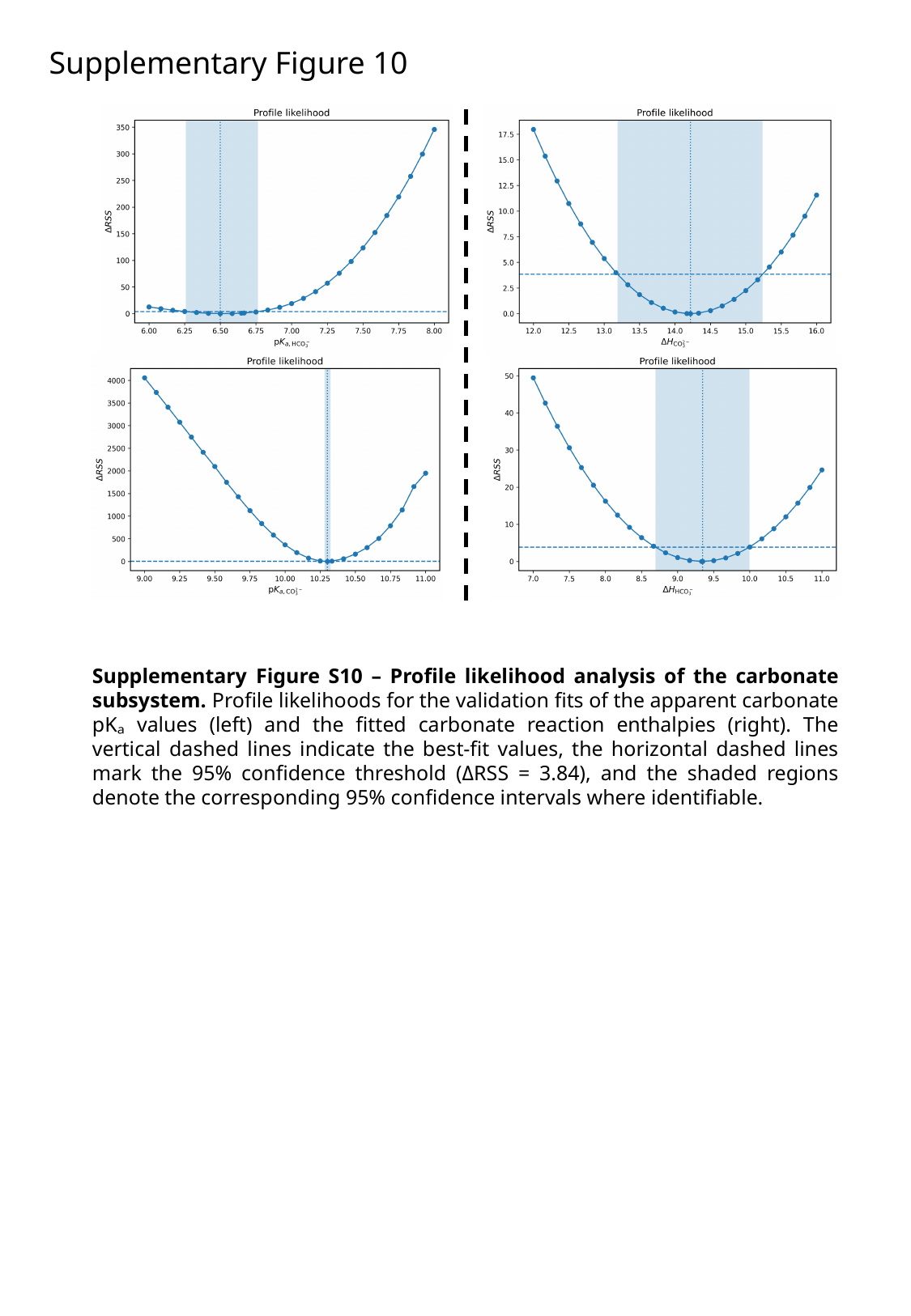

Supplementary Figure 10
Supplementary Figure S10 – Profile likelihood analysis of the carbonate subsystem. Profile likelihoods for the validation fits of the apparent carbonate pKₐ values (left) and the fitted carbonate reaction enthalpies (right). The vertical dashed lines indicate the best-fit values, the horizontal dashed lines mark the 95% confidence threshold (ΔRSS = 3.84), and the shaded regions denote the corresponding 95% confidence intervals where identifiable.

### Slide 11
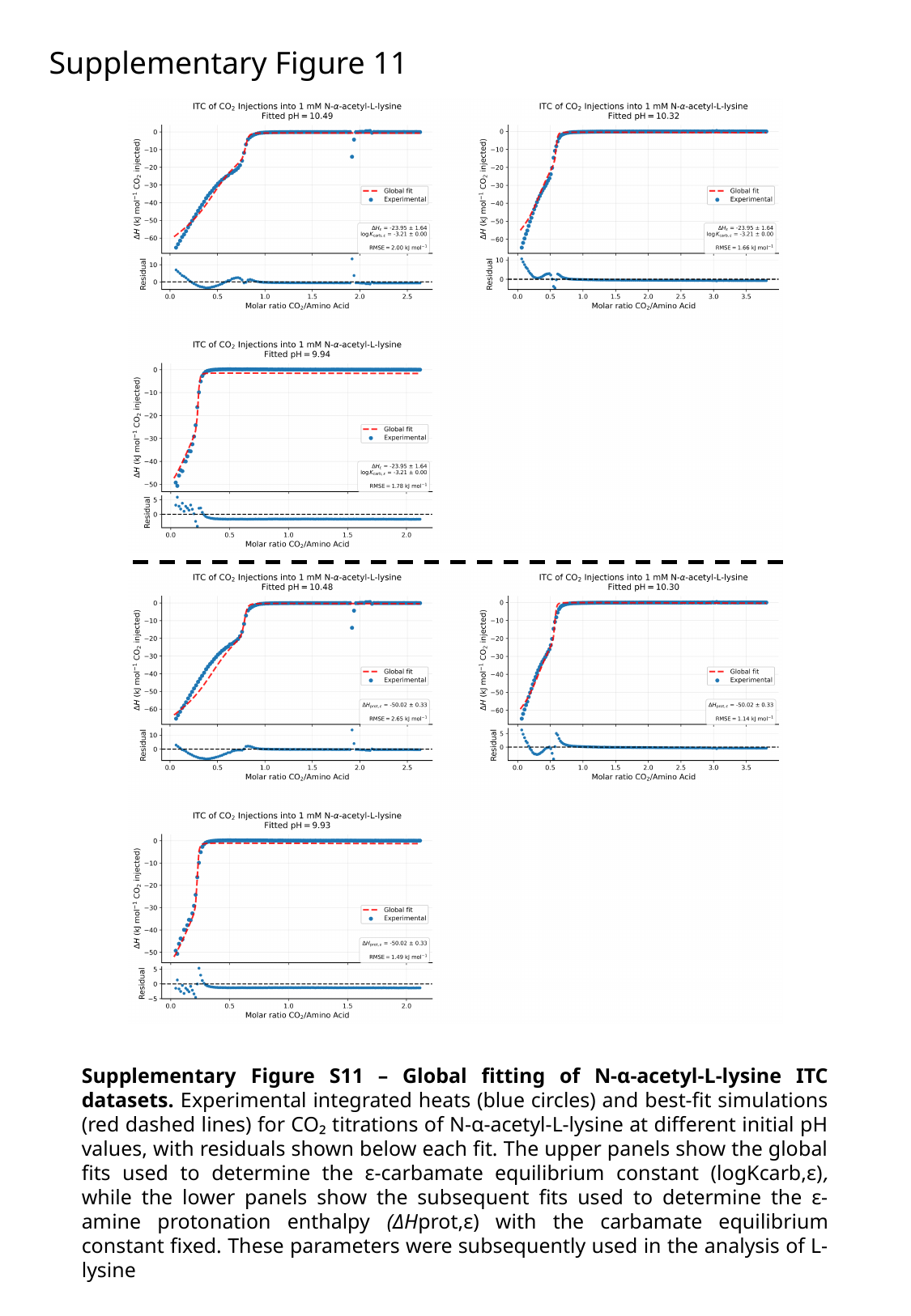

Supplementary Figure 11
Supplementary Figure S11 – Global fitting of N-α-acetyl-L-lysine ITC datasets. Experimental integrated heats (blue circles) and best-fit simulations (red dashed lines) for CO₂ titrations of N-α-acetyl-L-lysine at different initial pH values, with residuals shown below each fit. The upper panels show the global fits used to determine the ε-carbamate equilibrium constant (logKcarb,ε), while the lower panels show the subsequent fits used to determine the ε-amine protonation enthalpy (ΔHprot,ε) with the carbamate equilibrium constant fixed. These parameters were subsequently used in the analysis of L-lysine

### Slide 12
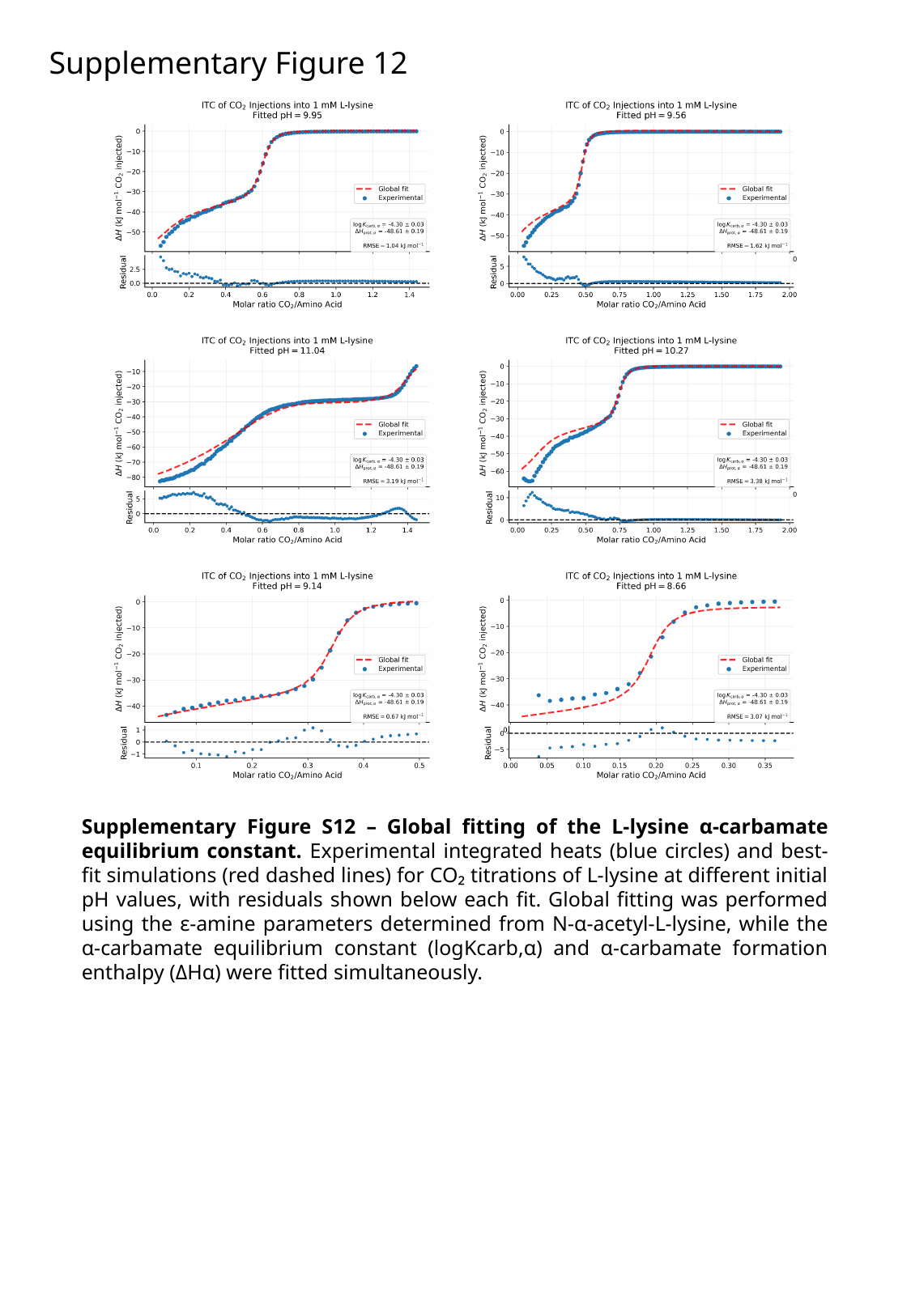

Supplementary Figure 12
Supplementary Figure S12 – Global fitting of the L-lysine α-carbamate equilibrium constant. Experimental integrated heats (blue circles) and best-fit simulations (red dashed lines) for CO₂ titrations of L-lysine at different initial pH values, with residuals shown below each fit. Global fitting was performed using the ε-amine parameters determined from N-α-acetyl-L-lysine, while the α-carbamate equilibrium constant (logKcarb,α) and α-carbamate formation enthalpy (ΔHα) were fitted simultaneously.

### Slide 13
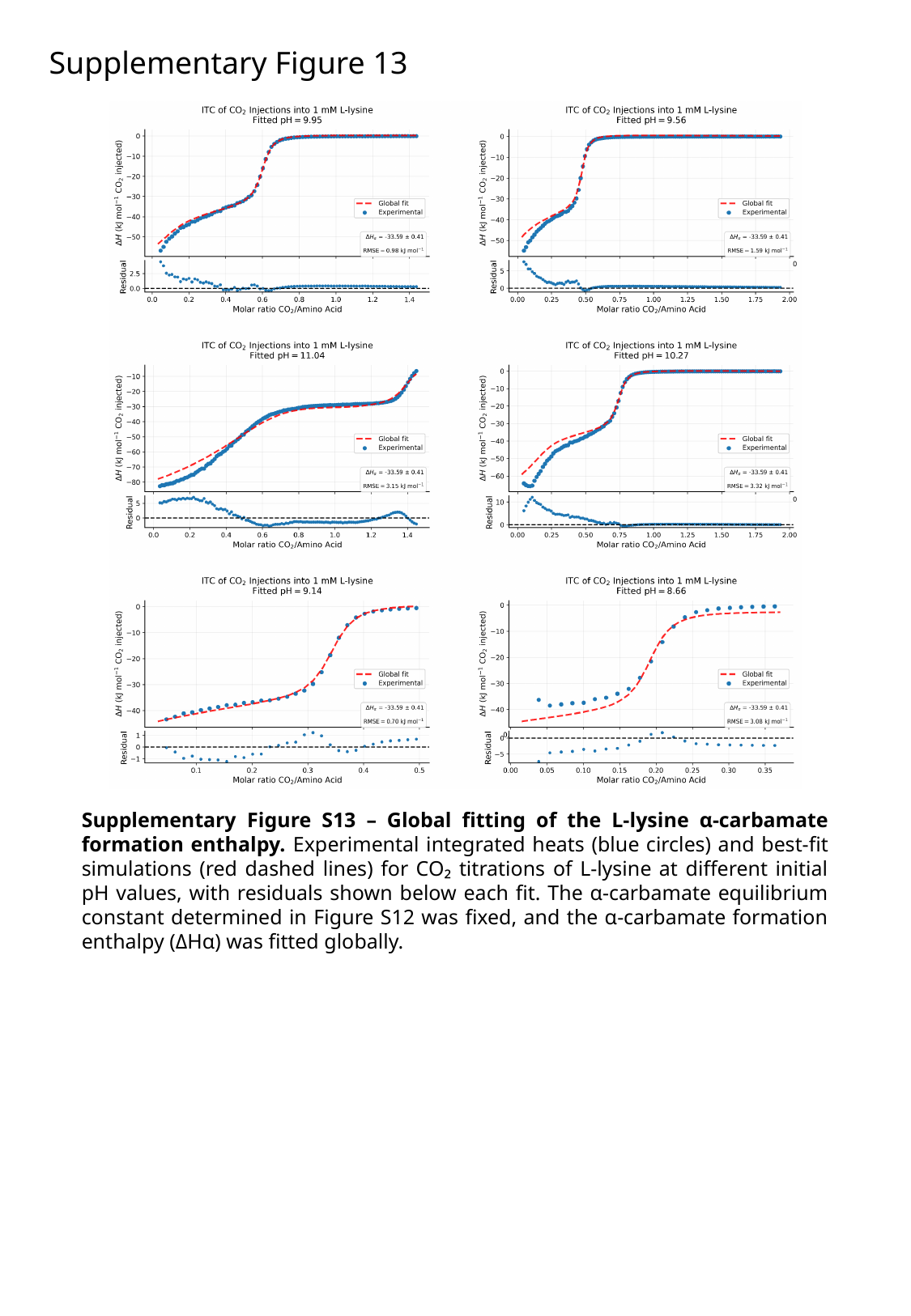

Supplementary Figure 13
Supplementary Figure S13 – Global fitting of the L-lysine α-carbamate formation enthalpy. Experimental integrated heats (blue circles) and best-fit simulations (red dashed lines) for CO₂ titrations of L-lysine at different initial pH values, with residuals shown below each fit. The α-carbamate equilibrium constant determined in Figure S12 was fixed, and the α-carbamate formation enthalpy (ΔHα) was fitted globally.

### Slide 14
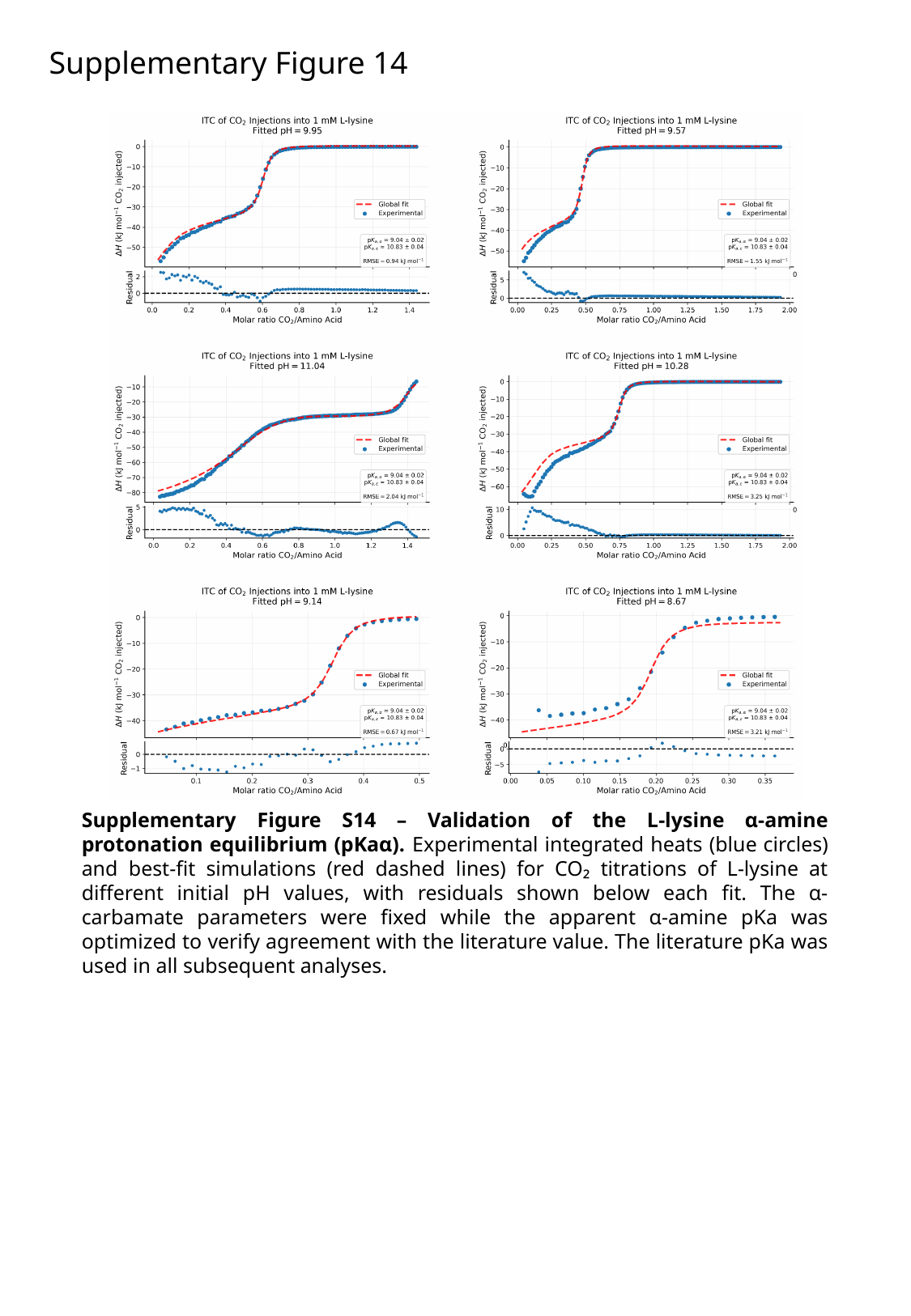

Supplementary Figure 14
Supplementary Figure S14 – Validation of the L-lysine α-amine protonation equilibrium (pKaα). Experimental integrated heats (blue circles) and best-fit simulations (red dashed lines) for CO₂ titrations of L-lysine at different initial pH values, with residuals shown below each fit. The α-carbamate parameters were fixed while the apparent α-amine pKa was optimized to verify agreement with the literature value. The literature pKa was used in all subsequent analyses.

### Slide 15
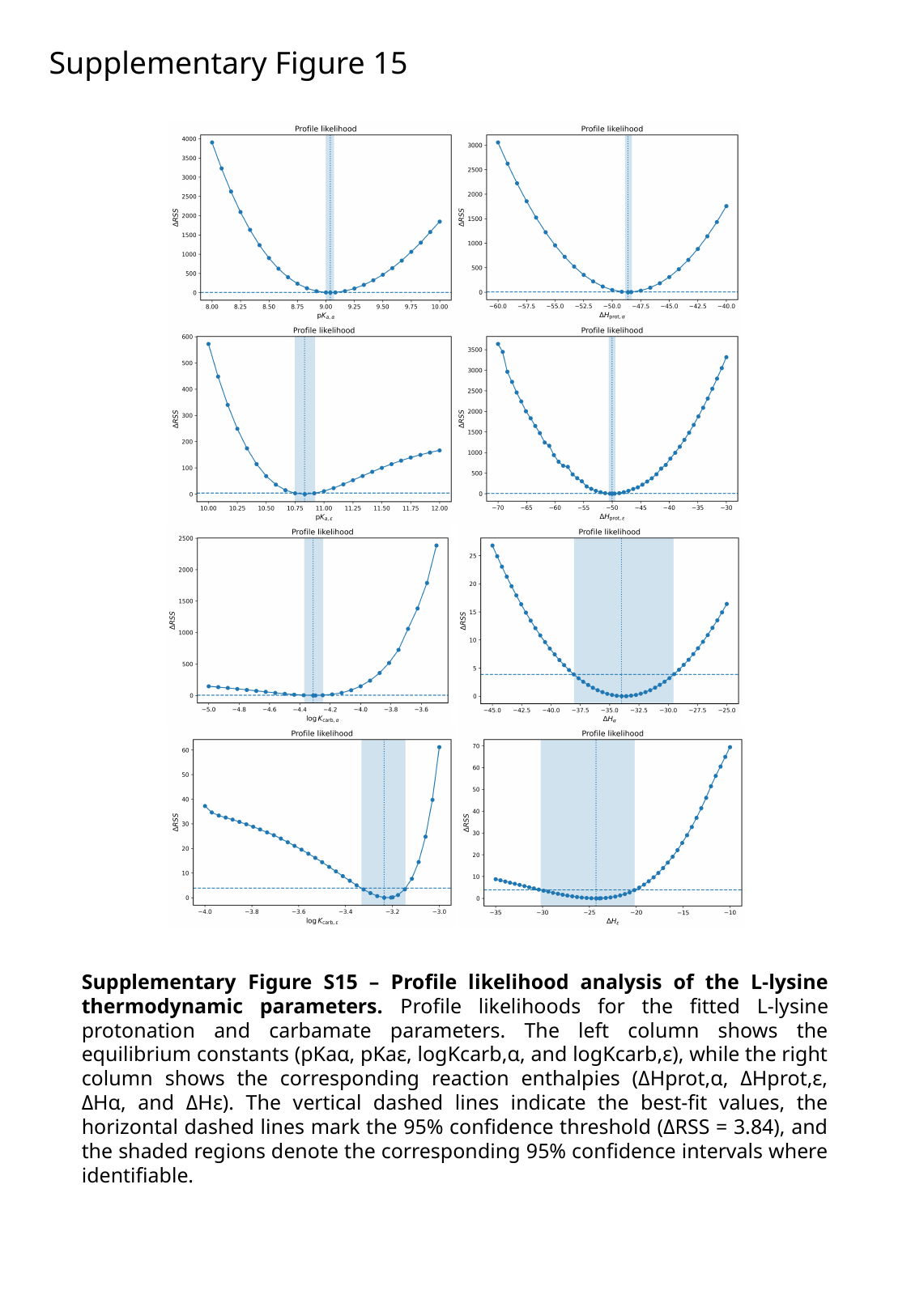

Supplementary Figure 15
Supplementary Figure S15 – Profile likelihood analysis of the L-lysine thermodynamic parameters. Profile likelihoods for the fitted L-lysine protonation and carbamate parameters. The left column shows the equilibrium constants (pKaα, pKaε, logKcarb,α, and logKcarb,ε), while the right column shows the corresponding reaction enthalpies (ΔHprot,α, ΔHprot,ε, ΔHα, and ΔHε). The vertical dashed lines indicate the best-fit values, the horizontal dashed lines mark the 95% confidence threshold (ΔRSS = 3.84), and the shaded regions denote the corresponding 95% confidence intervals where identifiable.

### Slide 16
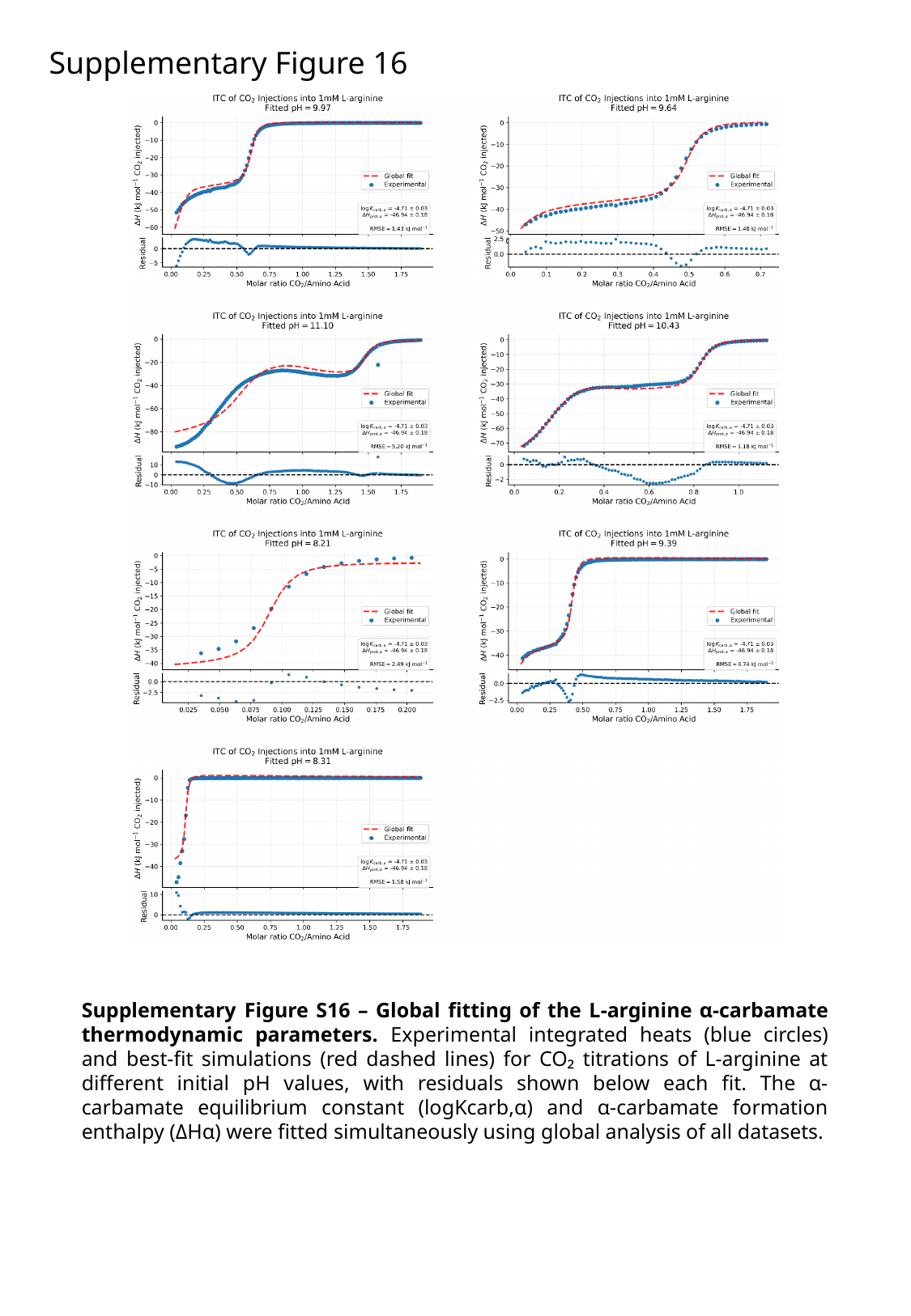

Supplementary Figure 16
Supplementary Figure S16 – Global fitting of the L-arginine α-carbamate thermodynamic parameters. Experimental integrated heats (blue circles) and best-fit simulations (red dashed lines) for CO₂ titrations of L-arginine at different initial pH values, with residuals shown below each fit. The α-carbamate equilibrium constant (logKcarb,α) and α-carbamate formation enthalpy (ΔHα) were fitted simultaneously using global analysis of all datasets.

### Slide 17
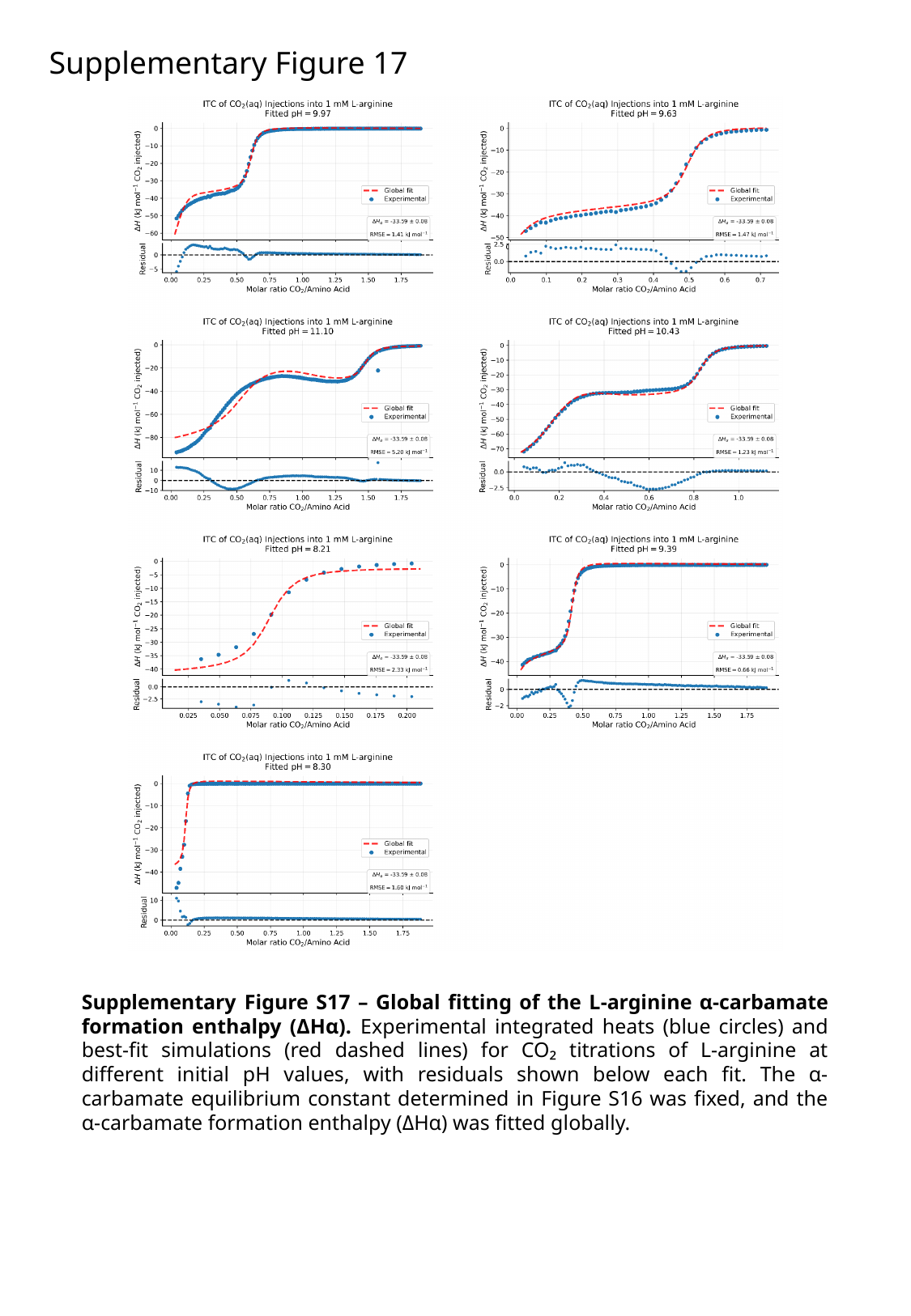

Supplementary Figure 17
Supplementary Figure S17 – Global fitting of the L-arginine α-carbamate formation enthalpy (ΔHα). Experimental integrated heats (blue circles) and best-fit simulations (red dashed lines) for CO₂ titrations of L-arginine at different initial pH values, with residuals shown below each fit. The α-carbamate equilibrium constant determined in Figure S16 was fixed, and the α-carbamate formation enthalpy (ΔHα) was fitted globally.

### Slide 18
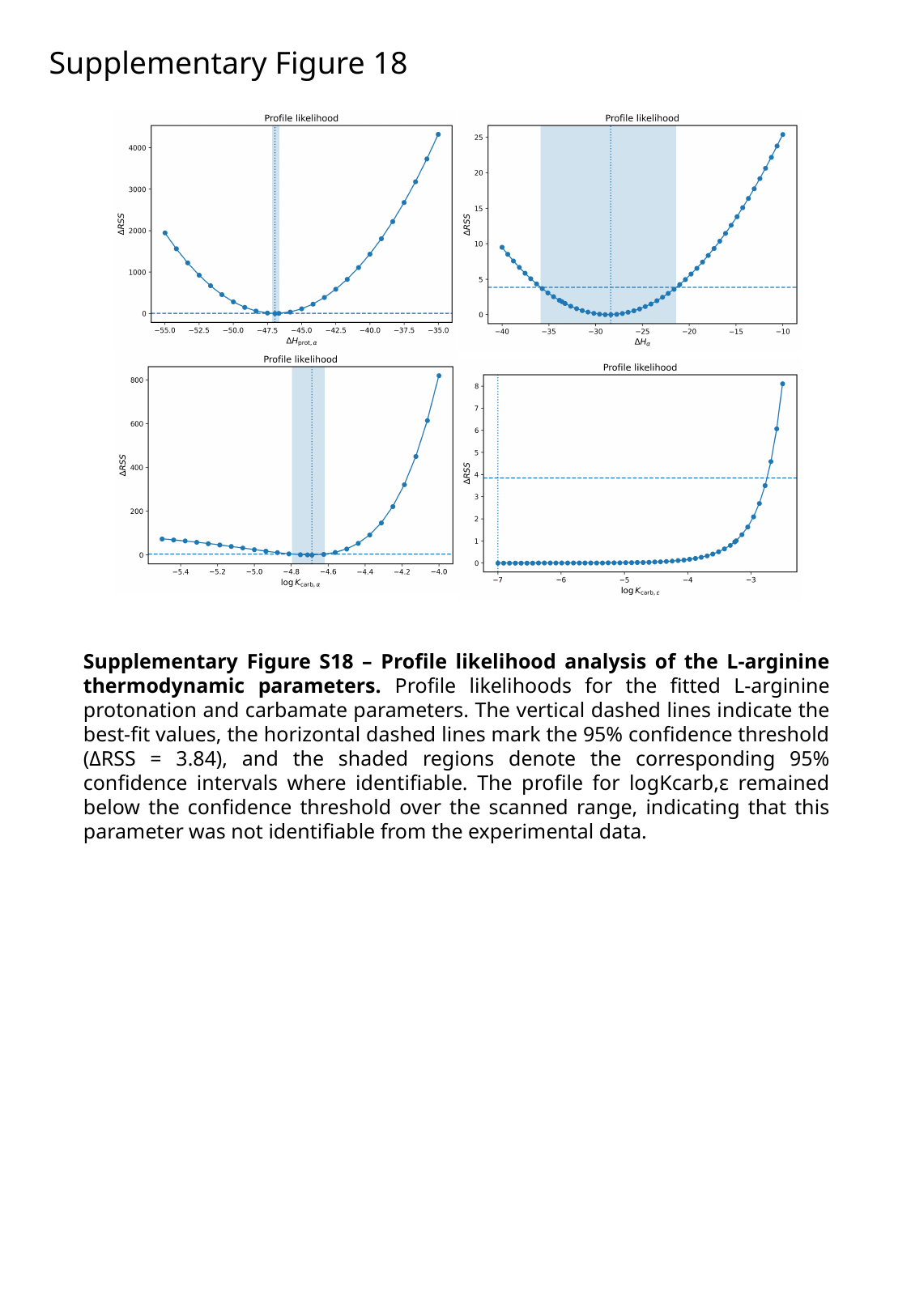

Supplementary Figure 18
Supplementary Figure S18 – Profile likelihood analysis of the L-arginine thermodynamic parameters. Profile likelihoods for the fitted L-arginine protonation and carbamate parameters. The vertical dashed lines indicate the best-fit values, the horizontal dashed lines mark the 95% confidence threshold (ΔRSS = 3.84), and the shaded regions denote the corresponding 95% confidence intervals where identifiable. The profile for logKcarb,ε remained below the confidence threshold over the scanned range, indicating that this parameter was not identifiable from the experimental data.

### Slide 19
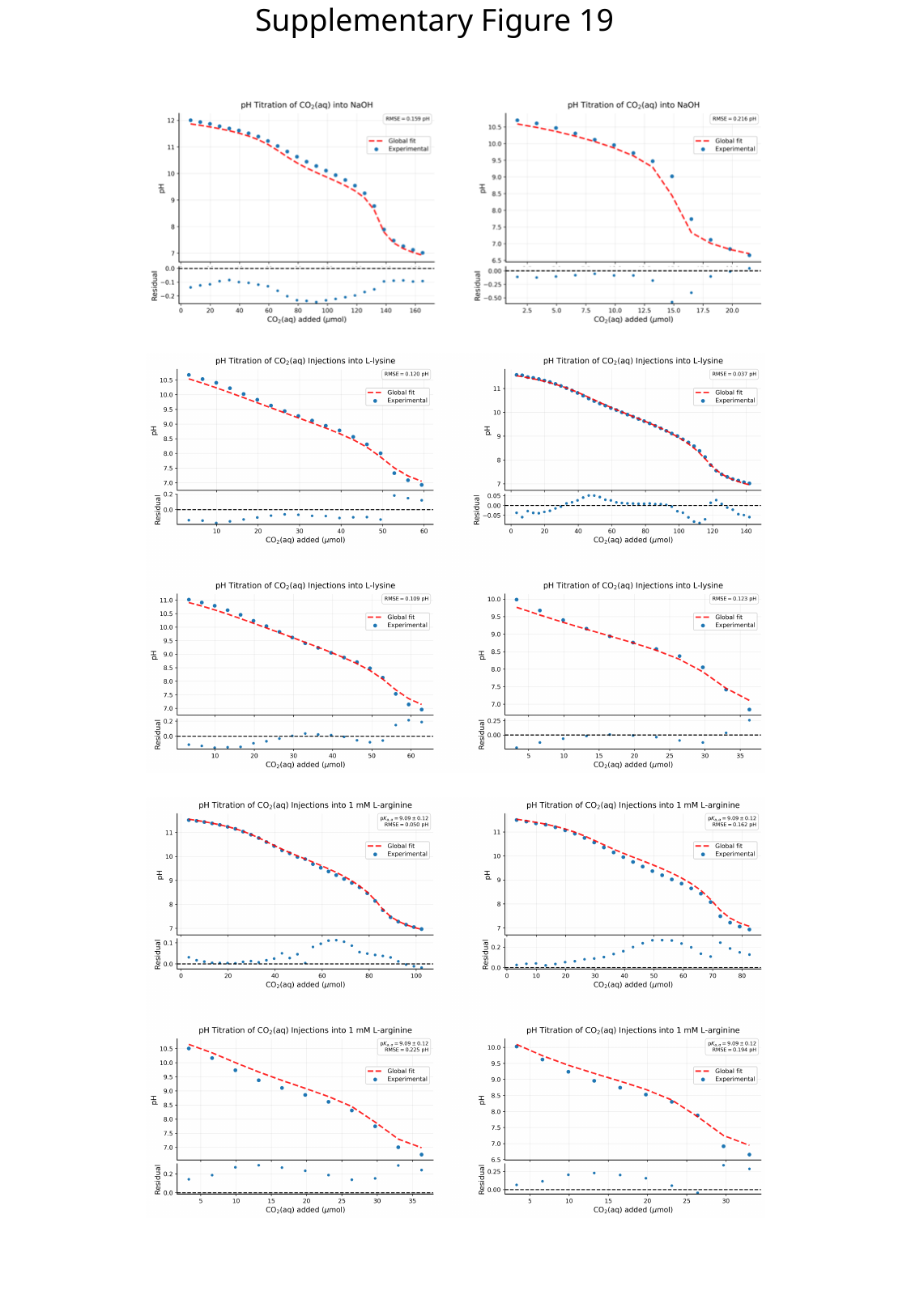

Supplementary Figure 19

### Slide 20
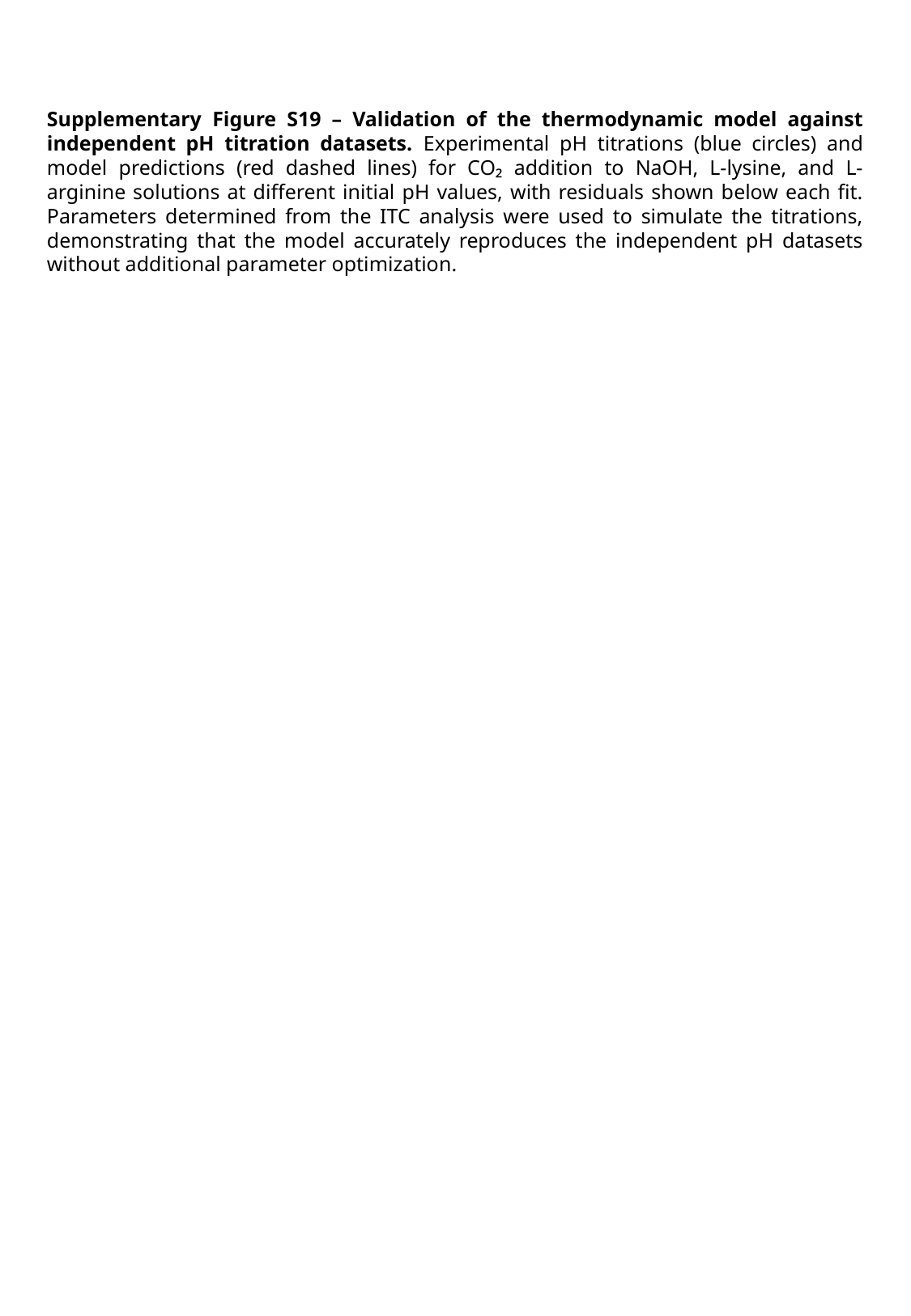

Supplementary Figure S19 – Validation of the thermodynamic model against independent pH titration datasets. Experimental pH titrations (blue circles) and model predictions (red dashed lines) for CO₂ addition to NaOH, L-lysine, and L-arginine solutions at different initial pH values, with residuals shown below each fit. Parameters determined from the ITC analysis were used to simulate the titrations, demonstrating that the model accurately reproduces the independent pH datasets without additional parameter optimization.

### Slide 21
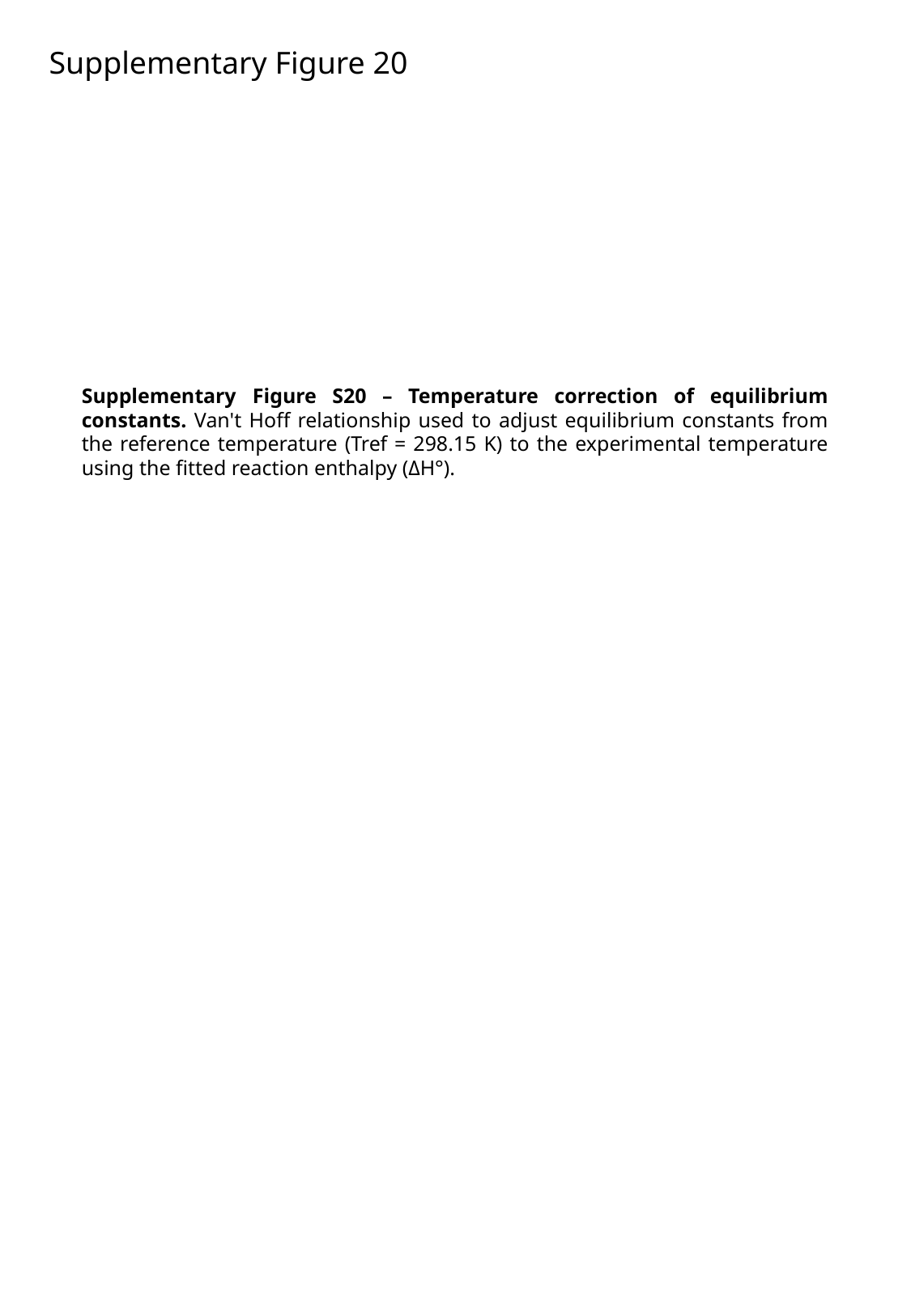

Supplementary Figure 20
Supplementary Figure S20 – Temperature correction of equilibrium constants. Van't Hoff relationship used to adjust equilibrium constants from the reference temperature (Tref = 298.15 K) to the experimental temperature using the fitted reaction enthalpy (ΔH°).

### Slide 22
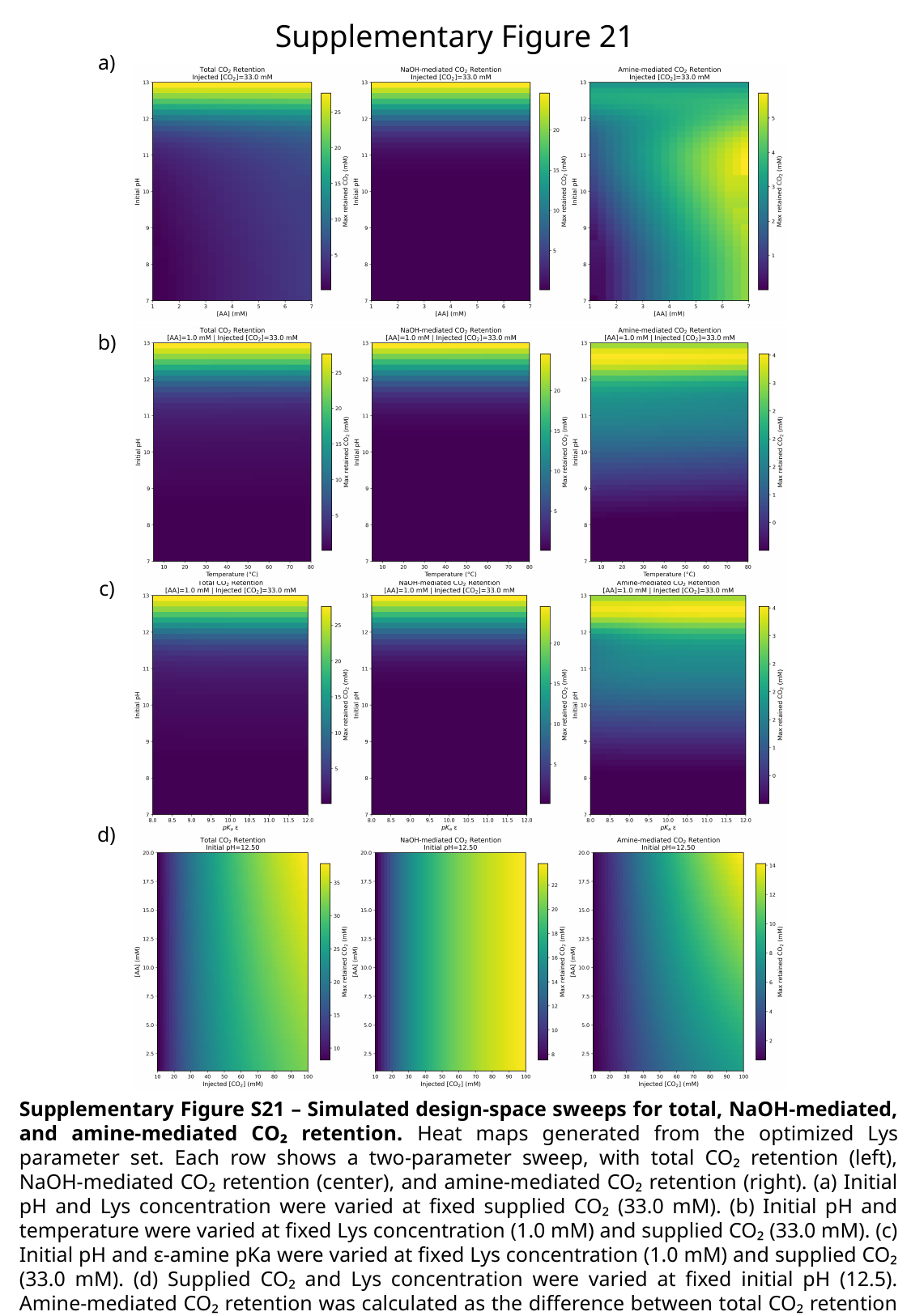

Supplementary Figure 21
a)
b)
c)
d)
Supplementary Figure S21 – Simulated design-space sweeps for total, NaOH-mediated, and amine-mediated CO₂ retention. Heat maps generated from the optimized Lys parameter set. Each row shows a two-parameter sweep, with total CO₂ retention (left), NaOH-mediated CO₂ retention (center), and amine-mediated CO₂ retention (right). (a) Initial pH and Lys concentration were varied at fixed supplied CO₂ (33.0 mM). (b) Initial pH and temperature were varied at fixed Lys concentration (1.0 mM) and supplied CO₂ (33.0 mM). (c) Initial pH and ε-amine pKa were varied at fixed Lys concentration (1.0 mM) and supplied CO₂ (33.0 mM). (d) Supplied CO₂ and Lys concentration were varied at fixed initial pH (12.5). Amine-mediated CO₂ retention was calculated as the difference between total CO₂ retention and the corresponding NaOH-only background.

### Slide 23
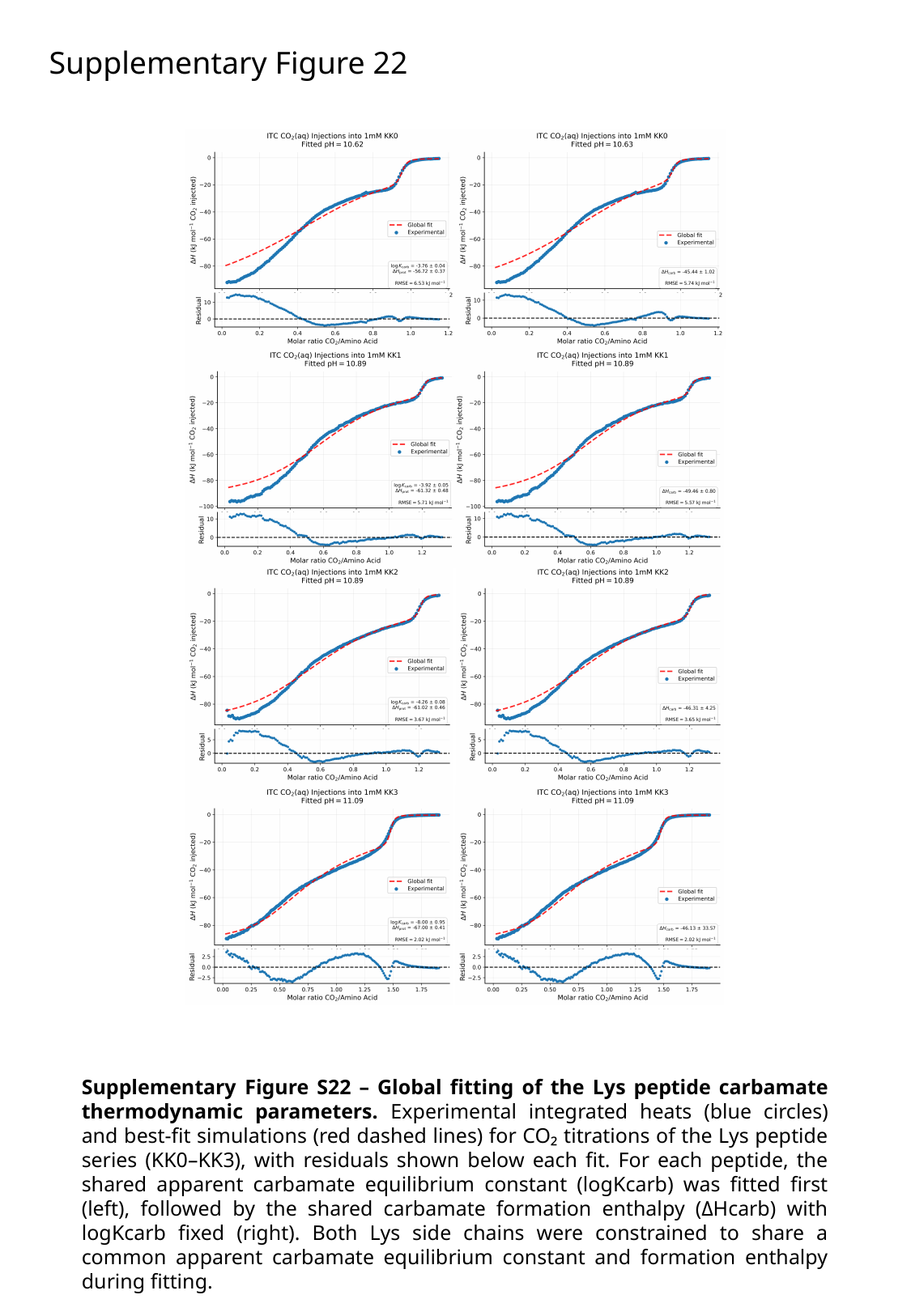

Supplementary Figure 22
Supplementary Figure S22 – Global fitting of the Lys peptide carbamate thermodynamic parameters. Experimental integrated heats (blue circles) and best-fit simulations (red dashed lines) for CO₂ titrations of the Lys peptide series (KK0–KK3), with residuals shown below each fit. For each peptide, the shared apparent carbamate equilibrium constant (logKcarb) was fitted first (left), followed by the shared carbamate formation enthalpy (ΔHcarb) with logKcarb fixed (right). Both Lys side chains were constrained to share a common apparent carbamate equilibrium constant and formation enthalpy during fitting.

### Slide 24
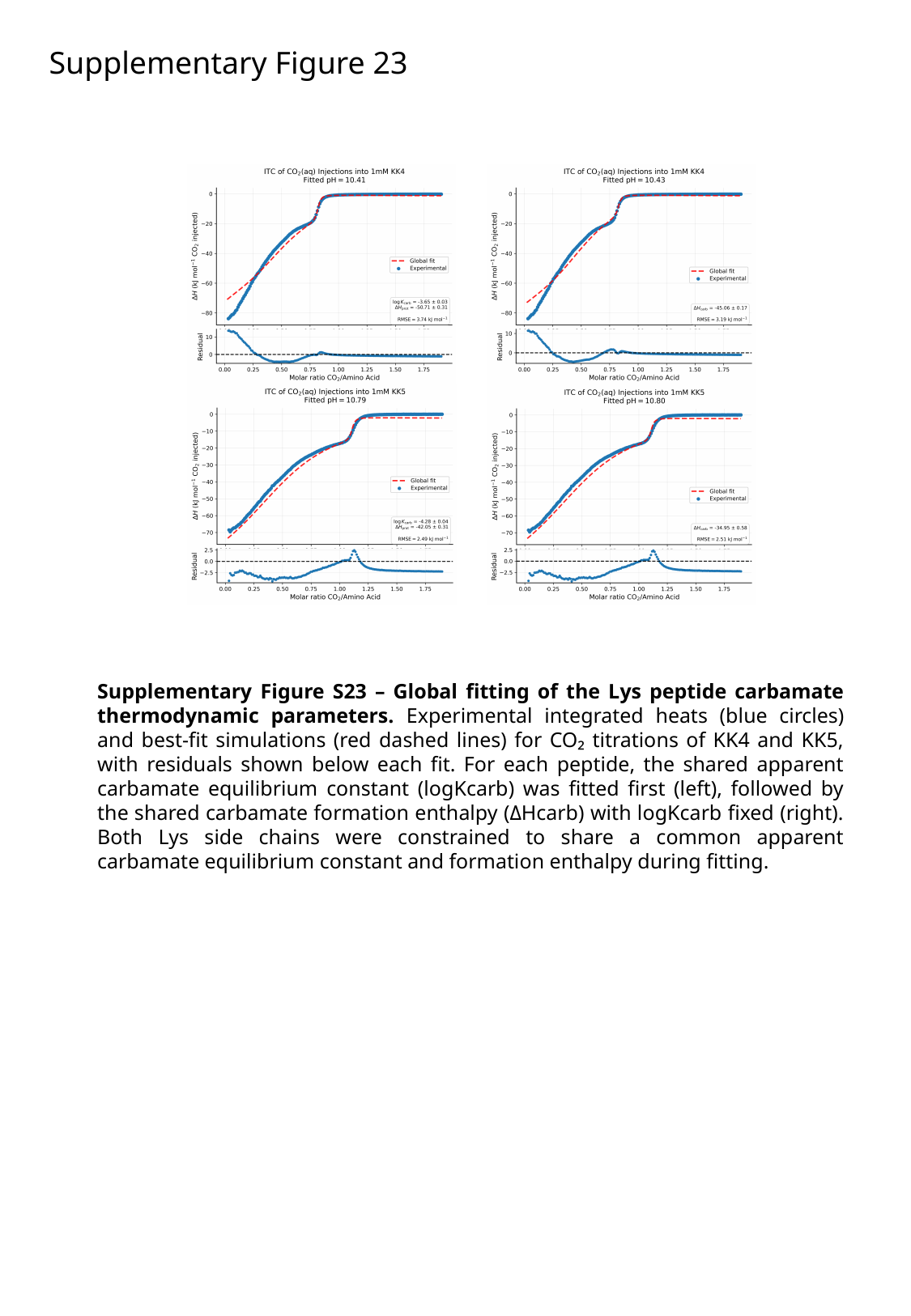

Supplementary Figure 23
Supplementary Figure S23 – Global fitting of the Lys peptide carbamate thermodynamic parameters. Experimental integrated heats (blue circles) and best-fit simulations (red dashed lines) for CO₂ titrations of KK4 and KK5, with residuals shown below each fit. For each peptide, the shared apparent carbamate equilibrium constant (logKcarb) was fitted first (left), followed by the shared carbamate formation enthalpy (ΔHcarb) with logKcarb fixed (right). Both Lys side chains were constrained to share a common apparent carbamate equilibrium constant and formation enthalpy during fitting.

### Slide 25
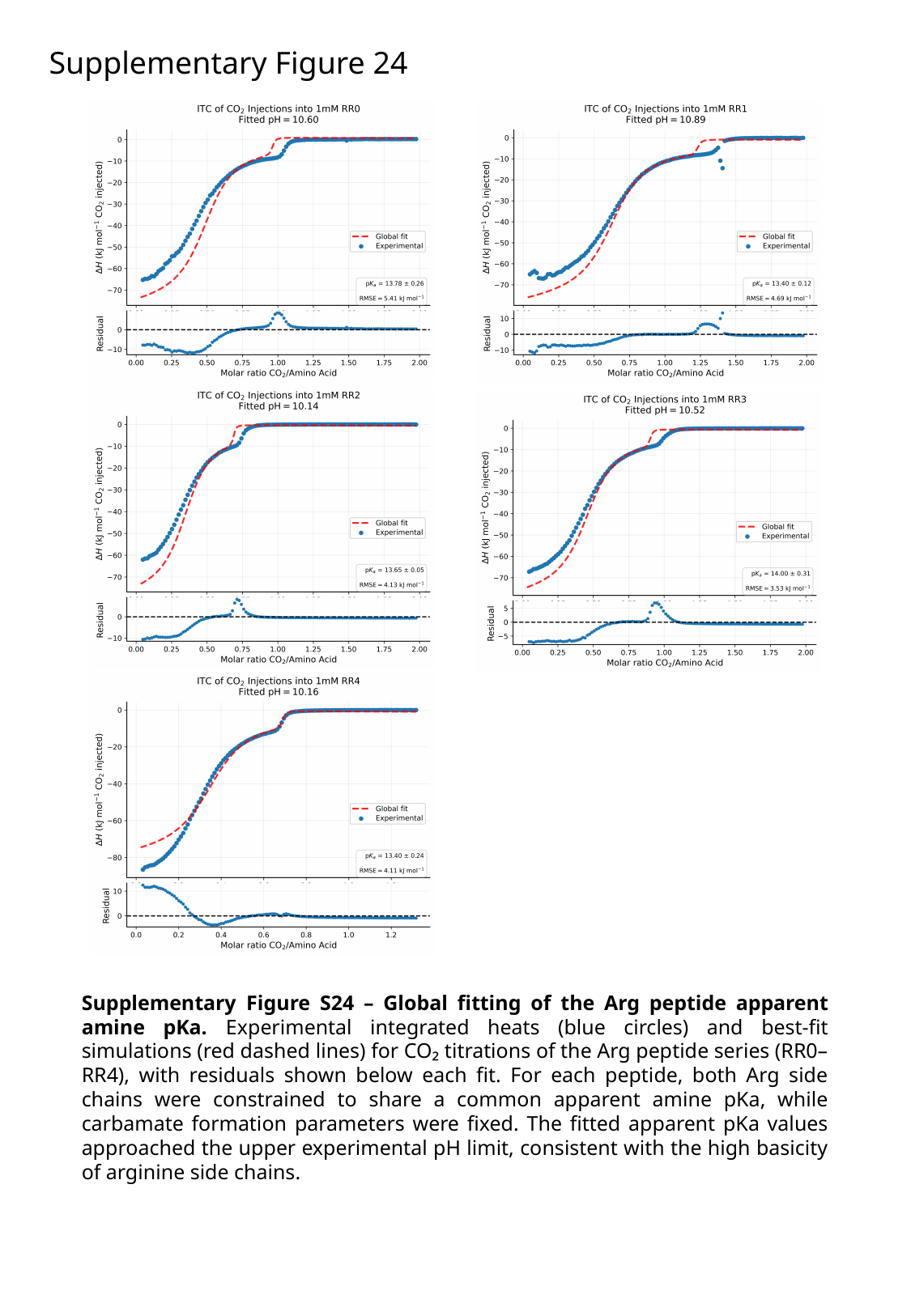

Supplementary Figure 24
Supplementary Figure S24 – Global fitting of the Arg peptide apparent amine pKa. Experimental integrated heats (blue circles) and best-fit simulations (red dashed lines) for CO₂ titrations of the Arg peptide series (RR0–RR4), with residuals shown below each fit. For each peptide, both Arg side chains were constrained to share a common apparent amine pKa, while carbamate formation parameters were fixed. The fitted apparent pKa values approached the upper experimental pH limit, consistent with the high basicity of arginine side chains.

### Slide 26
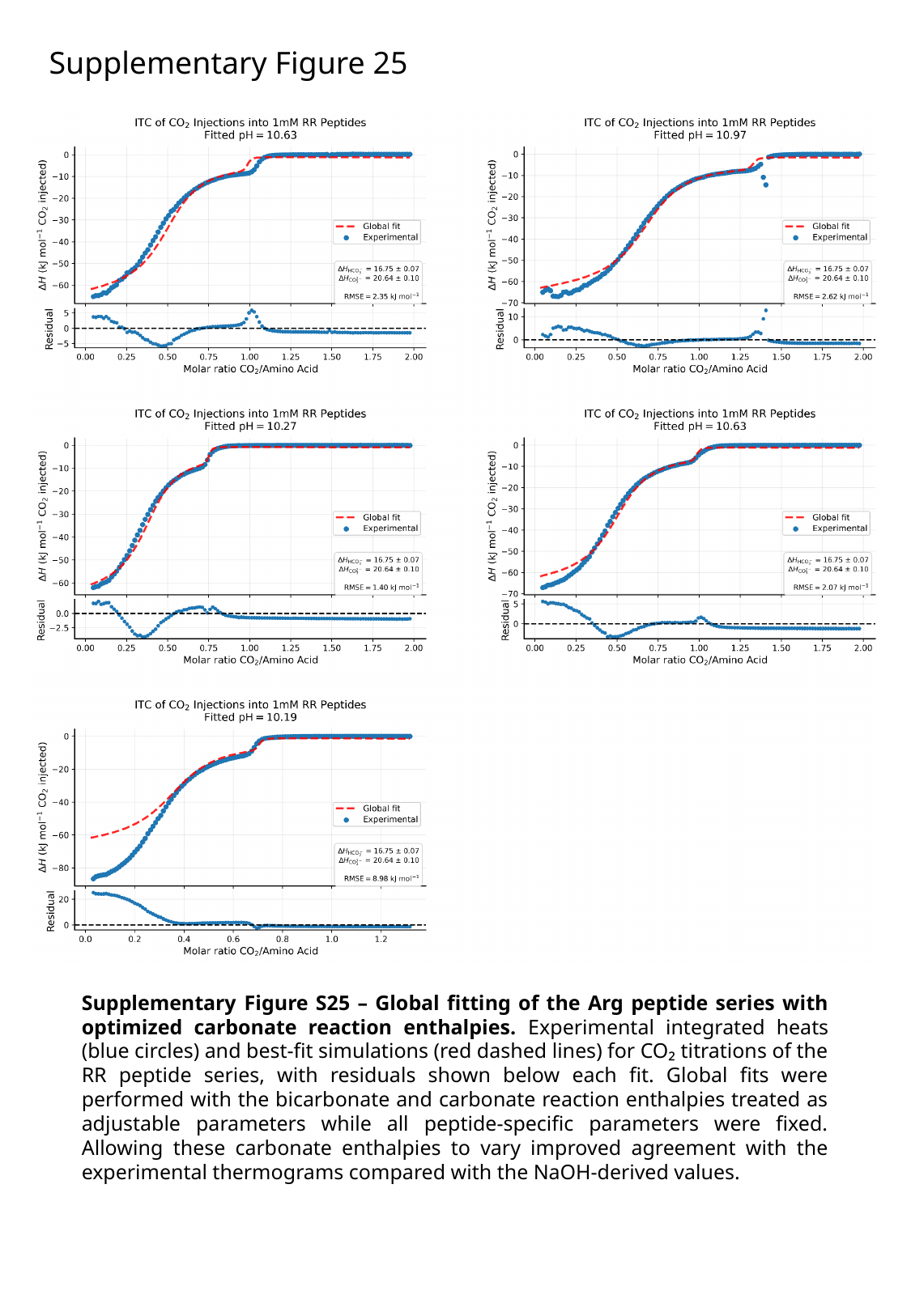

Supplementary Figure 25
Supplementary Figure S25 – Global fitting of the Arg peptide series with optimized carbonate reaction enthalpies. Experimental integrated heats (blue circles) and best-fit simulations (red dashed lines) for CO₂ titrations of the RR peptide series, with residuals shown below each fit. Global fits were performed with the bicarbonate and carbonate reaction enthalpies treated as adjustable parameters while all peptide-specific parameters were fixed. Allowing these carbonate enthalpies to vary improved agreement with the experimental thermograms compared with the NaOH-derived values.

### Slide 27
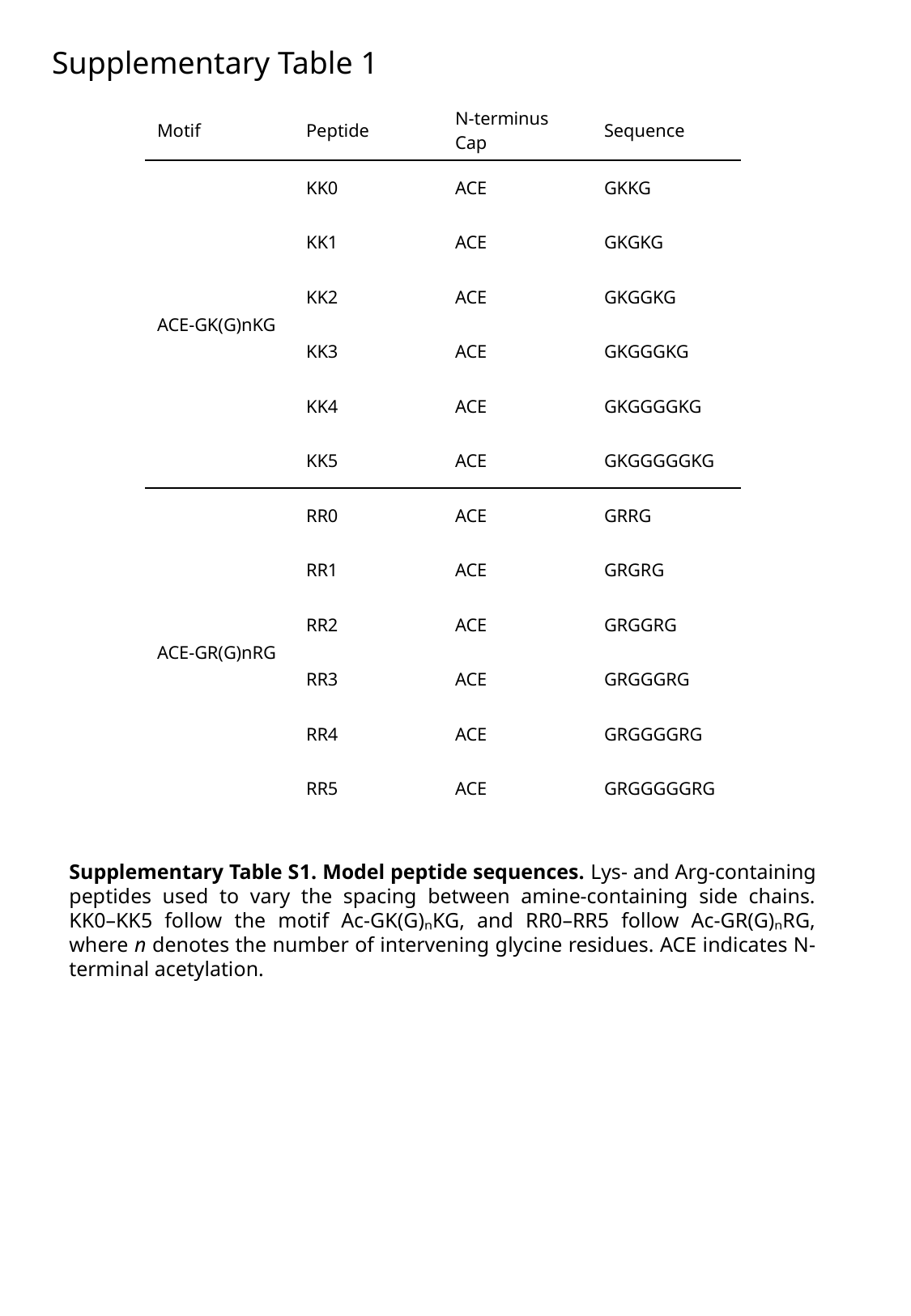

Supplementary Table 1
| Motif | Peptide | N-terminus Cap | Sequence |
| --- | --- | --- | --- |
| ACE-GK(G)nKG | KK0 | ACE | GKKG |
| | KK1 | ACE | GKGKG |
| | KK2 | ACE | GKGGKG |
| | KK3 | ACE | GKGGGKG |
| | KK4 | ACE | GKGGGGKG |
| | KK5 | ACE | GKGGGGGKG |
| ACE-GR(G)nRG | RR0 | ACE | GRRG |
| | RR1 | ACE | GRGRG |
| | RR2 | ACE | GRGGRG |
| | RR3 | ACE | GRGGGRG |
| | RR4 | ACE | GRGGGGRG |
| | RR5 | ACE | GRGGGGGRG |
Supplementary Table S1. Model peptide sequences. Lys- and Arg-containing peptides used to vary the spacing between amine-containing side chains. KK0–KK5 follow the motif Ac-GK(G)ₙKG, and RR0–RR5 follow Ac-GR(G)ₙRG, where n denotes the number of intervening glycine residues. ACE indicates N-terminal acetylation.

### Slide 28
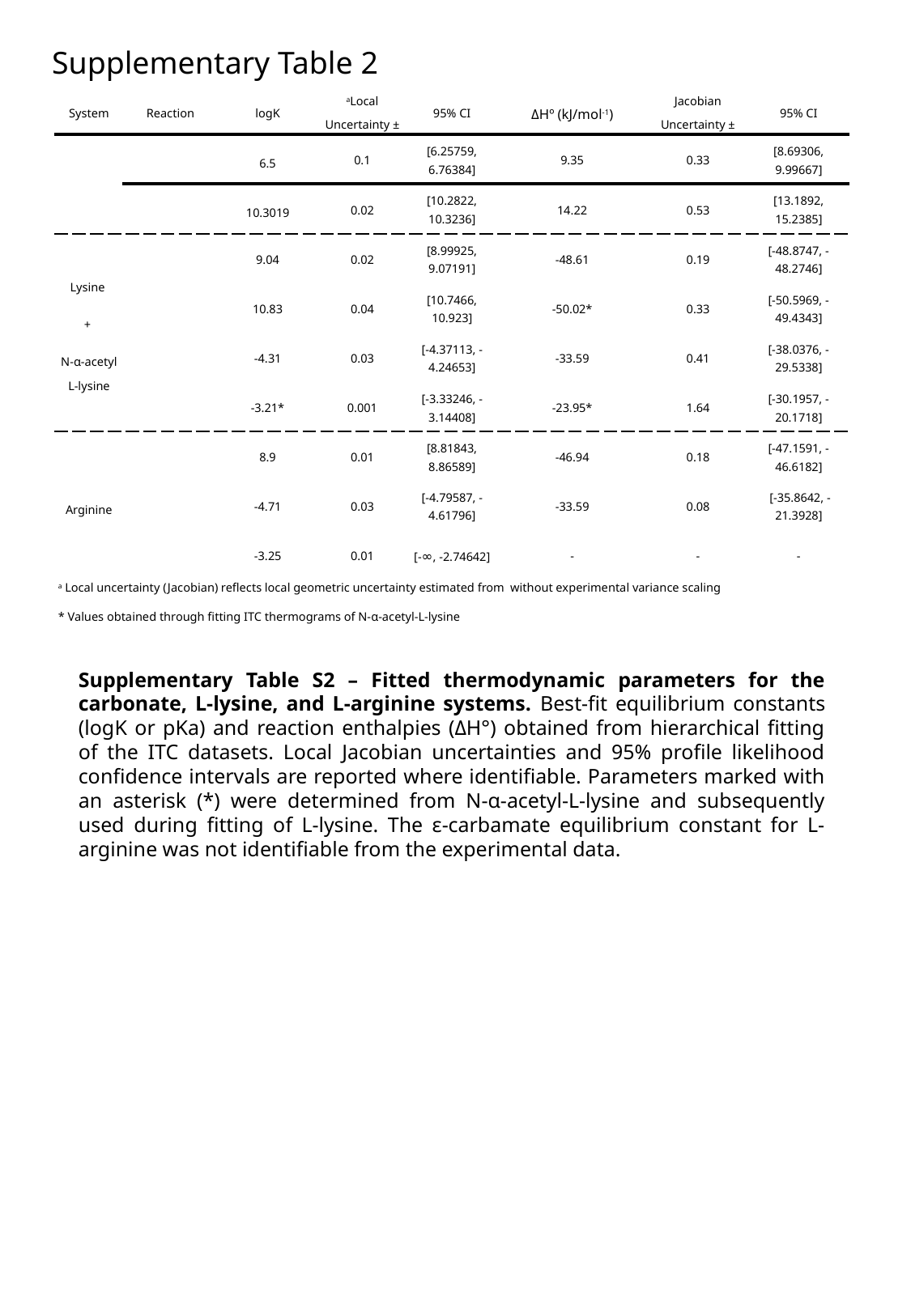

Supplementary Table 2
Supplementary Table S2 – Fitted thermodynamic parameters for the carbonate, L-lysine, and L-arginine systems. Best-fit equilibrium constants (logK or pKa) and reaction enthalpies (ΔH°) obtained from hierarchical fitting of the ITC datasets. Local Jacobian uncertainties and 95% profile likelihood confidence intervals are reported where identifiable. Parameters marked with an asterisk (*) were determined from N-α-acetyl-L-lysine and subsequently used during fitting of L-lysine. The ε-carbamate equilibrium constant for L-arginine was not identifiable from the experimental data.
